# CoalQC - Quality control while inferring demographic histories from genomic data: Application to forest tree genomes

**DOI:** 10.1101/2020.03.03.962365

**Authors:** Ajinkya Bharatraj Patil, Sagar Sharad Shinde, S Raghavendra, B.N Satish, C.G Kushalappa, Nagarjun Vijay

## Abstract

Estimating demographic histories using genomic datasets has proven to be useful in addressing diverse evolutionary questions. Despite improvements in inference methods and availability of large genomic datasets, quality control steps to be performed prior to the use of sequentially Markovian coalescent (SMC) based methods remains understudied. While various filtering and masking steps have been used by previous studies, the rationale for such filtering and its consequences have not been assessed systematically. In this study, we have developed a reusable pipeline called “CoalQC”, to investigate potential sources of bias (such as repeat regions, heterogeneous coverage, and callability). First, we demonstrate that genome assembly quality can affect the estimation of demographic history using the genomes of several species. We then use the CoalQC pipeline to evaluate how different repeat classes affect the inference of demographic history in the plant species *Populus trichocarpa.* Next, we assemble a draft genome by generating whole-genome sequencing data for *Mesua ferrea* (sampled from Western Ghats, India), a multipurpose forest plant distributed across tropical south-east Asia and use it as an example to evaluate several technical (sequencing technology, PSMC parameter settings) and biological aspects that need to be considered while comparing demographic histories. Finally, we collate the genomic datasets of 14 additional forest tree species to compare the temporal dynamics of Ne and find evidence of a strong bottleneck in all tropical forest plants during Mid-Pleistocene glaciations. Our findings suggest that quality control prior to the use of SMC based methods is important and needs to be standardised.

## Introduction

Coalescent theory continues to be the fundamental tool for the study of gene genealogies in a population genetic framework. Changes in coalescence time and the Watterson estimator of genetic diversity (θ_w_) along the genome serve as a record of the population history of a species through time. Increasing availability of whole genomic datasets for a multitude of species has made it possible to analyse demographic histories to answer a suite of questions such as host-parasite co-evolution (Hecht et al. 2018), effect of climate change on population dynamics (Bai et al. 2018), hybridization (Vijay et al. 2016), speciation events and split times between species (Cahill et al. 2016), history of inbreeding (Prado-Martinez et al. 2013), mutational meltdown (Rogers and Slatkin 2017), detecting population decline and addressing threats of extinction (Mays et al. 2018). Such widespread use of genomic datasets for coalescent inferences was made possible by the introduction of the PSMC method (Li and Durbin 2011) that requires only one diploid genome sequencing dataset. Technical advances in the use of genomic datasets for making demographic inferences and prevalence of multi-individual datasets facilitated by reductions in sequencing cost now allow integration of information across an increasing number of individuals (Schiffels and Durbin 2014; Terhorst et al. 2016; Palamara et al. 2018).

Despite the widespread use of demographic history inference methods like PSMC, many potential sources of bias due to data quality have been identified and efforts to reduce such effects are considered important. Earlier studies have shown that low coverage regions, ascertainment bias, hyperdiverse sequences, the fraction of usable data available as well as population structure will affect the estimation and interpretation of demographic histories (Li and Durbin 2011; Mazet et al. 2015; Nadachowska-Brzyska et al. 2016). Notably, some of the PSMC parameters or options such as mutation rate and generation time are known to drastically change the scaling of the curve, while the trajectory remains unchanged (Nadachowska-Brzyska et al. 2015). Detailed guidelines for the use of PSMC and MSMC are described elsewhere (Mather et al. 2020). Although the effect of genome assembly quality on demographic inferences has not been systematically assessed, it had been noted that genome quality could bias the results (Tiley et al. 2018). Intriguingly, a recent paper investigated the effect of genome quality and concluded that contemporary demographic inference methods are robust to the quality of the reference genome used (Patton et al. 2019).

In general practice, repetitive elements and low coverage regions of the genome are masked before the analyses to overcome biased inferences (Foote et al. 2016). It has been suggested that at least 75% or more of the genome should be retained after masking for robust inference of demographic history. However, organisms that have high (>30%) repeat content would violate either the masking criteria or the fraction of the genome to be retained. While several such quality control steps have been applied prior to the use of PSMC on genomic data, the rationale for performing specific filtering steps and the adverse consequences of skipping such quality control is understudied. A standardised quality control pipeline would be able to alleviate some of these challenges.

In this study, we have created a reusable pipeline (CoalQC) for evaluating the quality of genome-wide coalescence inferences and demonstrate the utility of the tool using a newly generated ∼180X coverage whole genome sequencing dataset of *Mesua ferrea* (a tropical tree species distributed across south-east Asia) as well as public re-sequencing datasets of several species. We investigate the relevance of various filtering practices and specifically answer the following questions:

1. Does genome assembly quality affect the inference of coalescence histories?
2. How does the coalescence history differ between repeat classes?
3. Which biological (change in genome size) and technical (sequencing platform used, PSMC parameter settings) factors influence demographic inference?
4. What can the comparison of demographic histories of forest plants reveal?

Our exploration of how several technical aspects can affect the inference of coalescence histories is relevant not just for use of the PSMC program, but also for numerous other tools that make coalescent inferences using genomic datasets. We also apply our pipeline to compare the demographic history of forest trees to evaluate whether ecologically relevant hypothesis can be robustly tested using demographic inference methods.

### New Approaches CoalQC

We have implemented a re-usable pipeline to perform quality control prior to the use of genome-wide coalescent methods. Separate modules to evaluate the effects of repeat regions, coverage and callability have been implemented to allow extensive quality control. The repeat module estimates independent demographic histories using genomic regions of one repeat class at a time along with non-repeat regions of the genome. Informative graphs that (a) compare these independent estimates of Ne, (b) quantify relative abundance of each repeat class in various atomic intervals, (c) assess robustness of the inferred results using bootstrap replicates, (d) visualise trends of change in heterozygosity and Ts/Tv ratio is generated by this module. We believe this module will be a valuable quality check prior to masking specific repeat regions of the genome.

The module for coverage is specifically designed to evaluate the robustness of the results to different coverage thresholds. Genomic regions are divided into several cumulative coverage classes based on the local read depth. These coverage classes are then used to independently estimate demographic histories and generate a comparative graph that can be used to understand the robustness of the results to coverage constraints. Similar to the coverage module, the callability module divides the genome into several callability classes to identify regions of the genome that need to be excluded from the analysis by masking. Detailed instructions and example commands for the use of the pipeline are provided on the github repository of the CoalQC program (https://github.com/ceglab/coalqc).

## Results

### Does genome quality affect demographic inference?

Genome assembly quality encompasses multiple factors such as sequence contiguity (generally quantified as N50), number and length of gaps, the fraction of genes assembled (quantified using BUSCO’s) and fraction of the genome assembled (quantified based on the percent of reads mapping to the genome assembly). To assess the effect of genome quality on demographic inference, we compared Ne trajectories estimated using a single human individual (NA12878) mapped to five different versions (hg4, hg10, hg15, hg19, and hg38) of human genome assemblies with varying levels of quality. We found that all the measures of genome quality used by us showed an improvement in recent versions of the human genome (see **Table S1**). The estimated effective population size (Ne) showed greater variability between genome assembly versions during ancient i.e., 1-7 MYA (Mean of standard deviations in Ne of each atomic interval from 43 to 64 = 0.84) and recent i.e., 0-15 KYA (Mean of standard deviations in Ne of each atomic interval from 0 to 6 = 0.82) compared to mid-time period i.e., 100-400 KYA (Mean of standard deviations in Ne of each atomic interval from 18 to 32 = 0.29) (see **Fig. 1**). PSMC trajectories of earlier (poorer assembly quality metrics) versions of the human genome showed higher estimates of Ne during the ancient (∼1-7 MYA) and recent times (∼0-15 KYA) and lower estimates of Ne during the mid-time period (∼100-400 KYA) compared to the recent versions of the human genome.

**Figure 1a:**
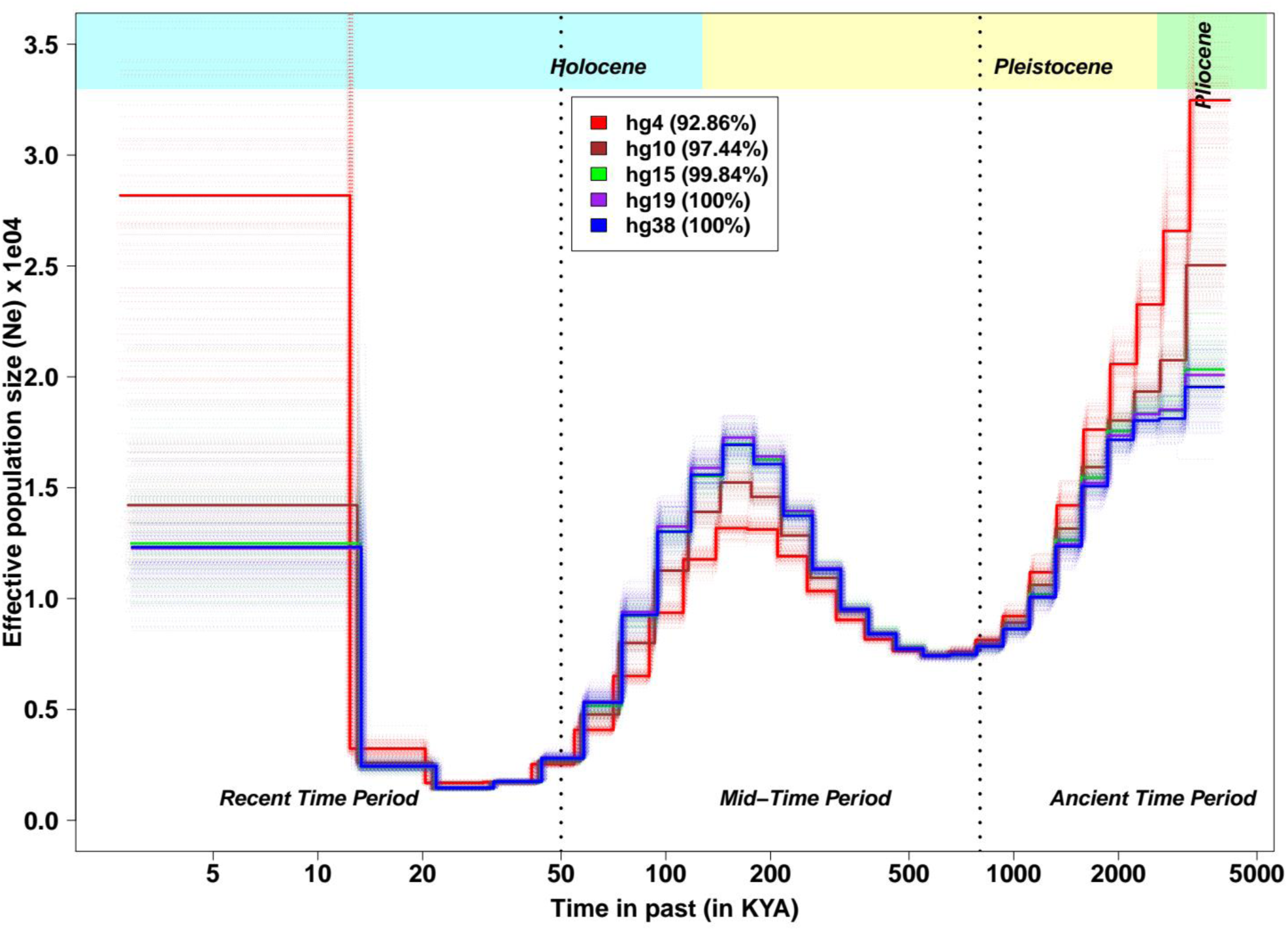
Change in Effective population size (N_e_) with change in the genome quality of Human assemblies. PSMC curve with bootstrap replicates for Human-NA12878 (see **Table S2**) mapped to human assembly version 4 (hg4) shown in red, to hg10 shown in brown, to hg15 shown in green, to hg19 shown in purple and to hg38 shown in blue, corresponding mapping percentages are given in parentheses. Poor quality assembly (hg4) overestimated the Ne during recent (∼1 KYA) and ancient (∼1-5 MYA) times, whereas Ne was underestimated during mid-period (∼100-400 KYA) compared to better assemblies.

To evaluate the effect of assembly quality on the robustness of results we performed 100 bootstrap runs using each of the human genome versions considered. The heterogeneity between bootstrap replicates within each version was quantified as the coefficient of variation (CV) across the 100 replicates. While the CV of the early version of human genome assembly (Mean of the CV of Ne across bootstrap replicates of hg4 assembly=0.054) was higher than that of the other recent assemblies (hg10=0.041, hg15=0.042, hg19=0.045, and hg38=0.043) considered by us, the CV’s of all the assemblies are very similar (see **Fig. 1b**). Comparable estimates of Ne across bootstrap runs suggest that these estimates are being robustly inferred for each specific genome assembly.

**Figure 1b:**
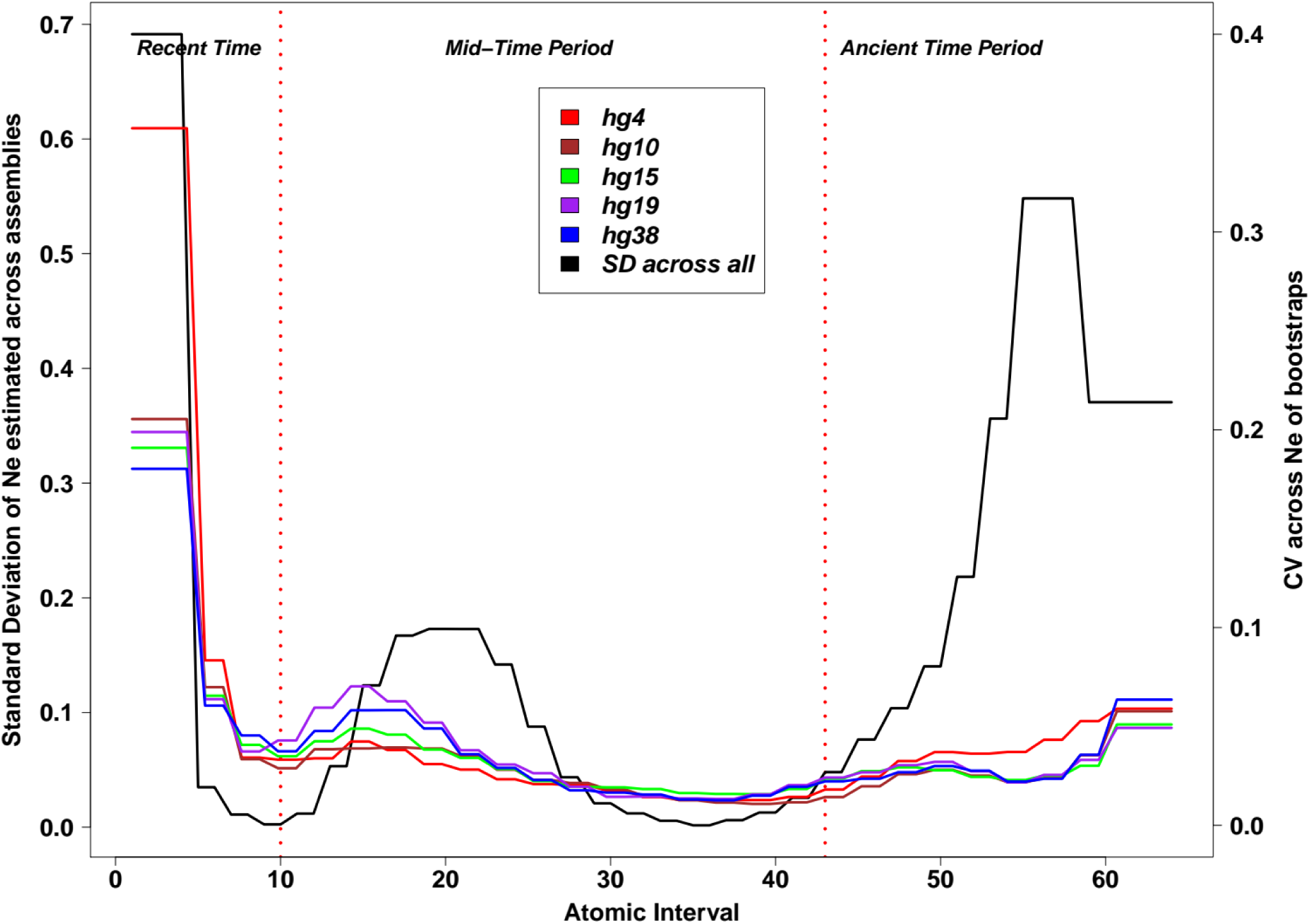
Extent of variation in PSMC trajectories inferred from Human assemblies. Estimates of Ne inferred using each assembly showed heterogeneity across time points. Blackline indicates Standard Deviation in each atomic interval across estimates of all the assemblies. The colored lines show the Coefficient of Variation within 100 bootstrap replicates of each assembly. SD curve (black line) shows that estimates in the Atomic intervals contributing to recent times (AI 1-6) and ancient times (AI 43-64) showed the highest variation across assemblies. The CV of the poorest assembly (hg4) shows the highest variation across bootstrap estimates in recent and ancient times suggesting relatively low robustness compared to others but there was not much difference for mid-period.

To ensure that the effect of genome assembly quality is not limited to just the human genome, we compared the demographic histories inferred from the initial and recent versions of the *Tribolium castaneum* and *Danio rerio* genomes (see **Table S1**). We find that similar to the differences seen between different versions of the human genome assembly, the estimates of Ne inferred from different versions of the genome show distinct trends (see **Fig. S1**). Our results from the human, red flour beetle (*Tribolium castaneum*) and zebrafish (*Danio rerio*) genomes suggest that genome quality does have a noticeable effect on demographic inference.

### How do repeat regions affect demographic inference?

Prior to performing demographic inference, repeat regions of the genome are generally masked and excluded from the analysis. Masking of repeat regions is justified by the high risk of assembly errors, collapsed segmental duplications and miss-mapping of short-reads in repeat regions. Plant genomes with a high fraction of repetitive content are more prone to be affected by repeats. Hence, we decided to use the high-quality genome of the plant *Populus trichocarpa* to compare the Ne trajectories inferred using masked and unmasked genomes to understand the magnitude of the change introduced by masking of repeat regions. The estimates of Ne from the masked compared to the unmasked genome were lower during ancient time period i.e., after ∼ 1 MYA (Mean difference in Ne across atomic intervals 48 to 64=0.63 x 10^4^) and higher during recent times i.e., 20KYA – 100 KYA (Mean difference in Ne across atomic intervals 5 to 17=0.45 x 10^4^, see **Fig. 2a**).

**Figure 2a:**
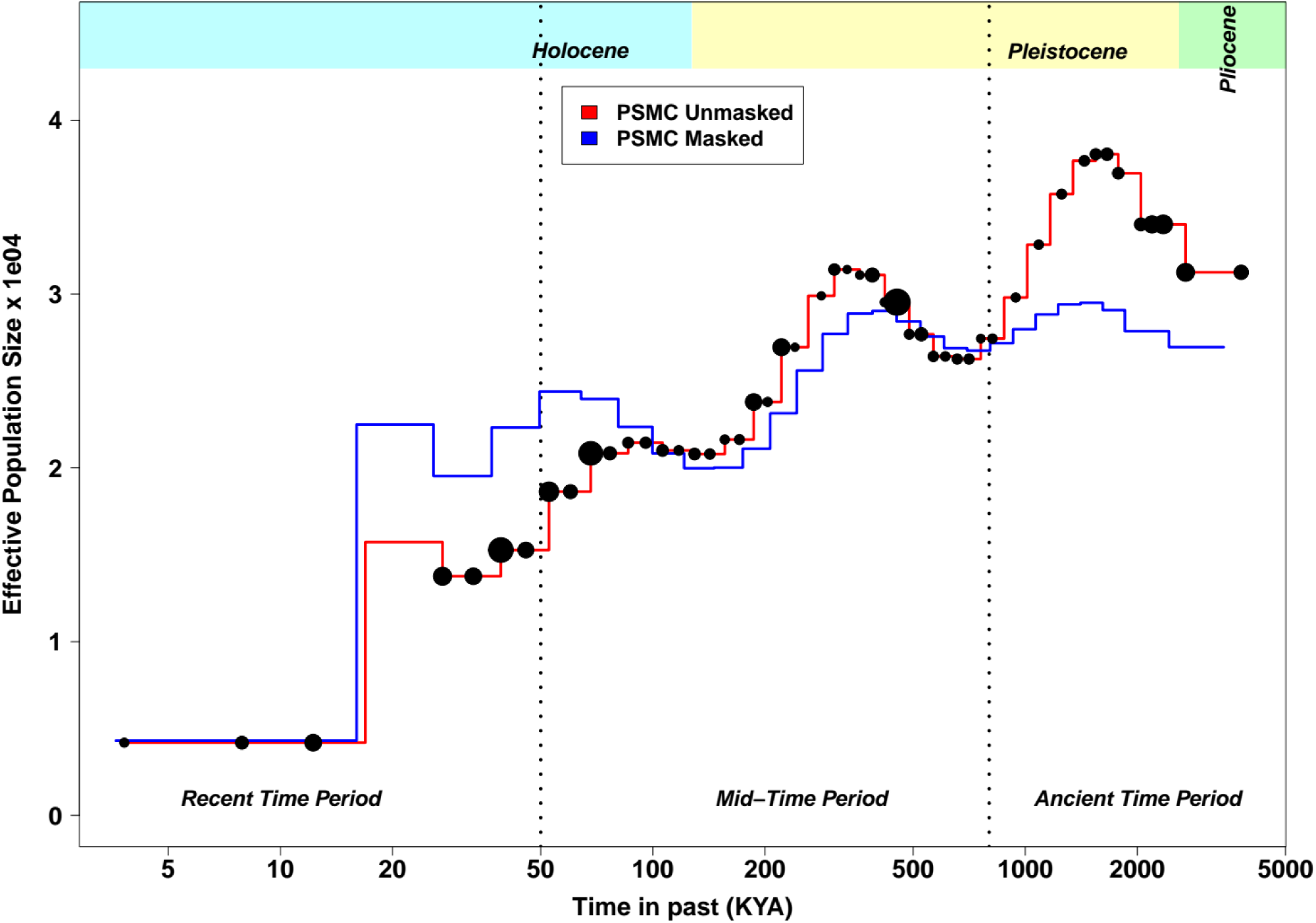
Change in Effective population Sizes (N_e_) due to masking of repeat regions in Populus trichocarpa *PSMC*. PSMC curve for *Populus trichocarpa* after masking all the repeat regions in the genome (blue line) and without masking (orange-red line). Unmasked trajectory has dots indicating the fraction of repeats in an atomic interval, larger the size more the repeat content in an atomic interval. Ne estimates during recent (20-100 KYA) and ancient (1MYA-5MYA) times show considerable differences between the two curves showing the effect of exclusion/inclusion of repeat sequences.

To evaluate how specific repeat classes affect the estimates of Ne, each repeat family was unmasked while keeping other repeats masked (see **Fig. 2b**). Estimates of Ne after inclusion of LTR-Gypsy was intermediate between masked and unmasked genome-based inferences during the ancient past i.e., after 1MYA (Mean difference in Ne compared to masked genome across atomic intervals 48 to 64= 0.29 x 10^4^), whereas it showed a similar trend as the unmasked genome during recent times i.e., 20KYA-100KYA (Mean difference in Ne compared to masked genome across atomic intervals 5 to 17=0.44 x 10^4^; see **Fig. 2c**). Other repeat classes did not influence the estimates as much as LTRs and were closer to the masked inference (see **Fig. 2b**). The robustness of the Ne estimates was assessed based on the variability (quantified as CV) between bootstrap replicates using the non-repeat fraction of the genome along with each individual repeat class. The CV was heterogeneous between repeat classes and was relatively higher in recent time intervals (see **Fig. 2c**). Robustness of the estimated values of Ne was comparable between the unmasked (Mean of the CV across atomic intervals= 0.04687) and masked (Mean of the CV across atomic intervals= 0.047) genomes.

**Figure 2b:**
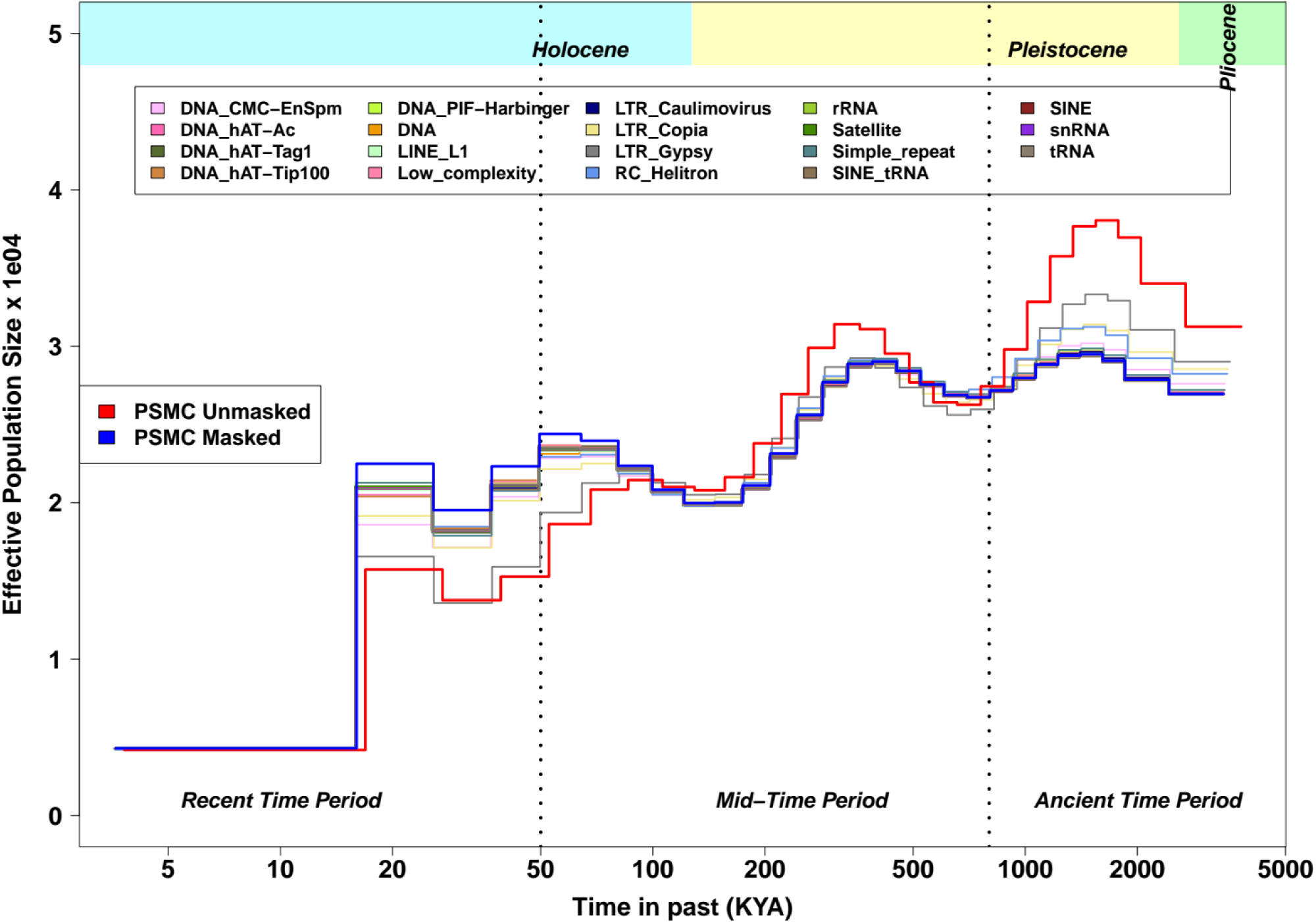
Change in Effective population Sizes (N_e_) after the inclusion of each repeat class in *Populus trichocarpa* PSMC separately. PSMC curves for *Populus trichocarpa,* with masked (blue) and unmasked (orange-red) genomes used. Change in trajectory due to the inclusion of each class of repeat to the masked genome is shown. Including each repeat class and masking, other repeat-classes will show changes specific to respective repeat-class. The inclusion of LTR-Gypsy shows a distinguishingly different trajectory similar to the unmasked genome during ∼20-100 KYA, which shows that the inclusion of LTR-Gypsy is influencing the trajectory in recent times.

**Figure 2c:**
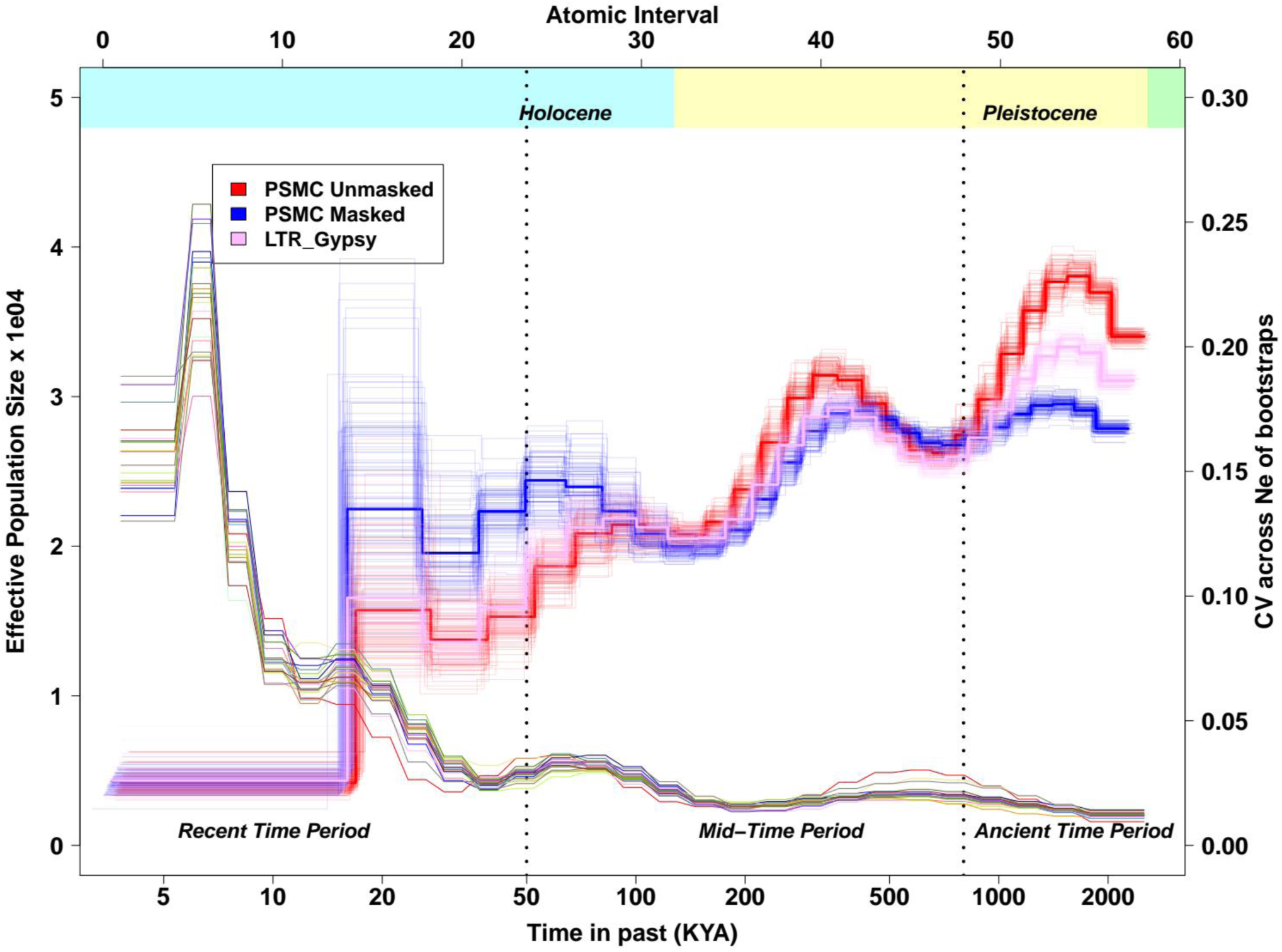
Bootstrapped PSMC results after masking of repeats in *Populus trichocarpa*. PSMC curves for *Populus trichocarpa* showing the robustness of changes due to masking of repeats. Masked (blue) and unmasked (orange-red) shows completely distinctive trajectories whereas unmasking only LTR-Gypsy repeat class (pink) also shows a marked difference. The second y-axis (red) shows the Coefficient of variation (CV) across the bootstraps across all the repeat classes. This indicates changes in Ne due to repeats are robust to bootstrap replications.

We found that the fraction of repeat content in a particular atomic interval was positively correlated (τ = 0.346, p-value= 0.0003, see **Fig. S2**) with the absolute difference between masked and unmasked genome-based estimates of effective population size (Ne). Having established that greater repeat abundance would more strongly affect estimates of Ne, we quantified repeat family-wise abundance in genomic regions corresponding to each atomic interval. In all atomic intervals, the non-repeat fraction was found to be the most abundant (see **Fig. 2d**). Among the repeat classes, LTR-Gypsy had the highest abundance in most of the atomic intervals. The extremely high abundance of LTR-Gypsy repeats in the first few atomic intervals could have led to the drastic change in the Ne trajectory during recent times (i.e., 20KYA-100KYA) after inclusion of LTR-Gypsy repeats. LTR’s and RC-Helitron have high abundance at the genome-wide level (see **Fig. 2e**) and have a greater influence on the estimates of Ne (see **Fig. 2b**).

**Figure 2d:**
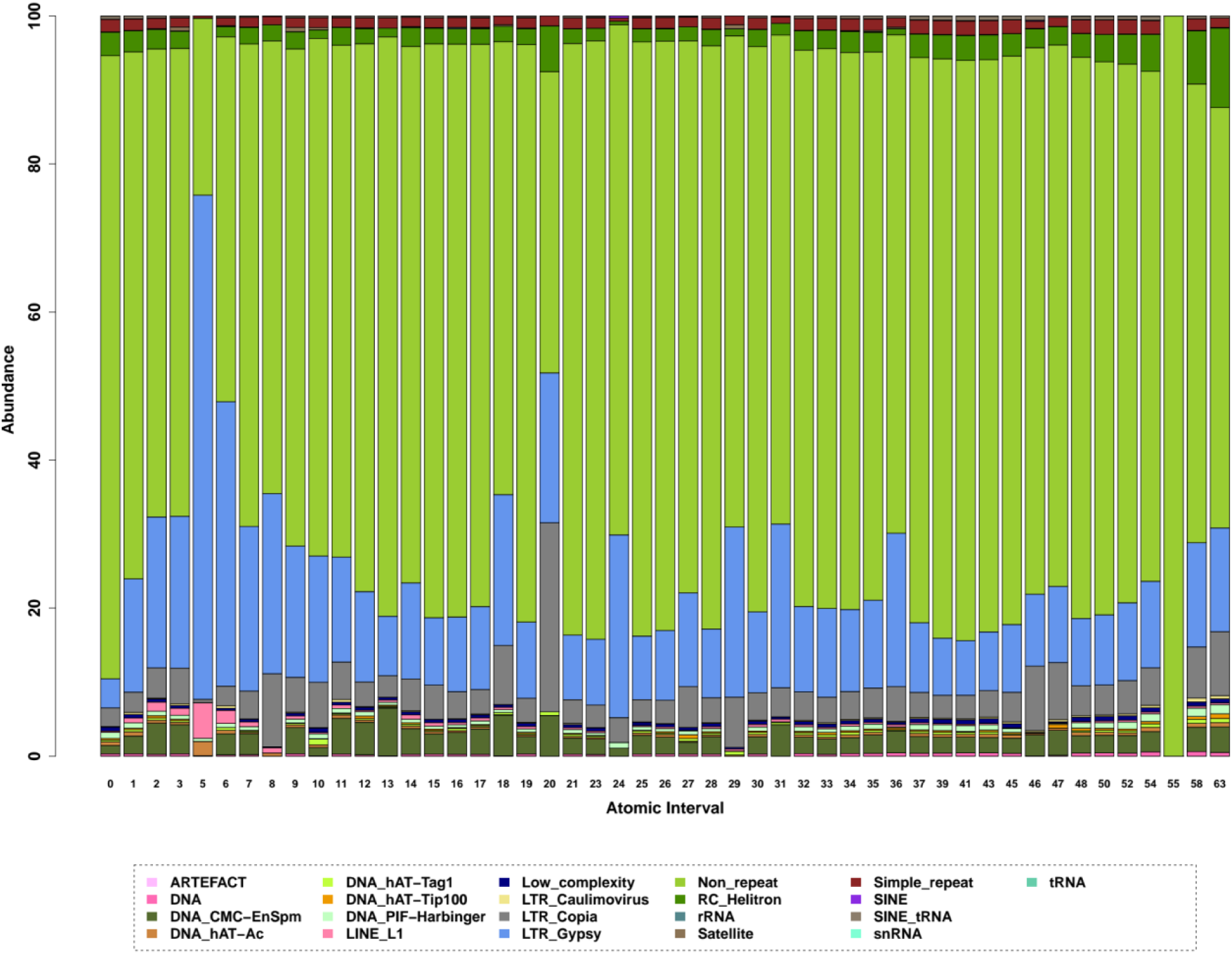
Abundance of different repeat classes across all atomic intervals in *Populus trichocarpa* PSMC. Contribution of various repeat classes to each atomic interval is shown. Non-repeat regions (light green) are generally most abundant across all intervals whereas some repeat classes such as LTR’s have considerable abundance in some of the atomic intervals. Repeat families such as LTR-Gypsy, LTR-Copia and RC-Helitron showed higher abundance compared to other repeat classes.

**Figure 2e:**
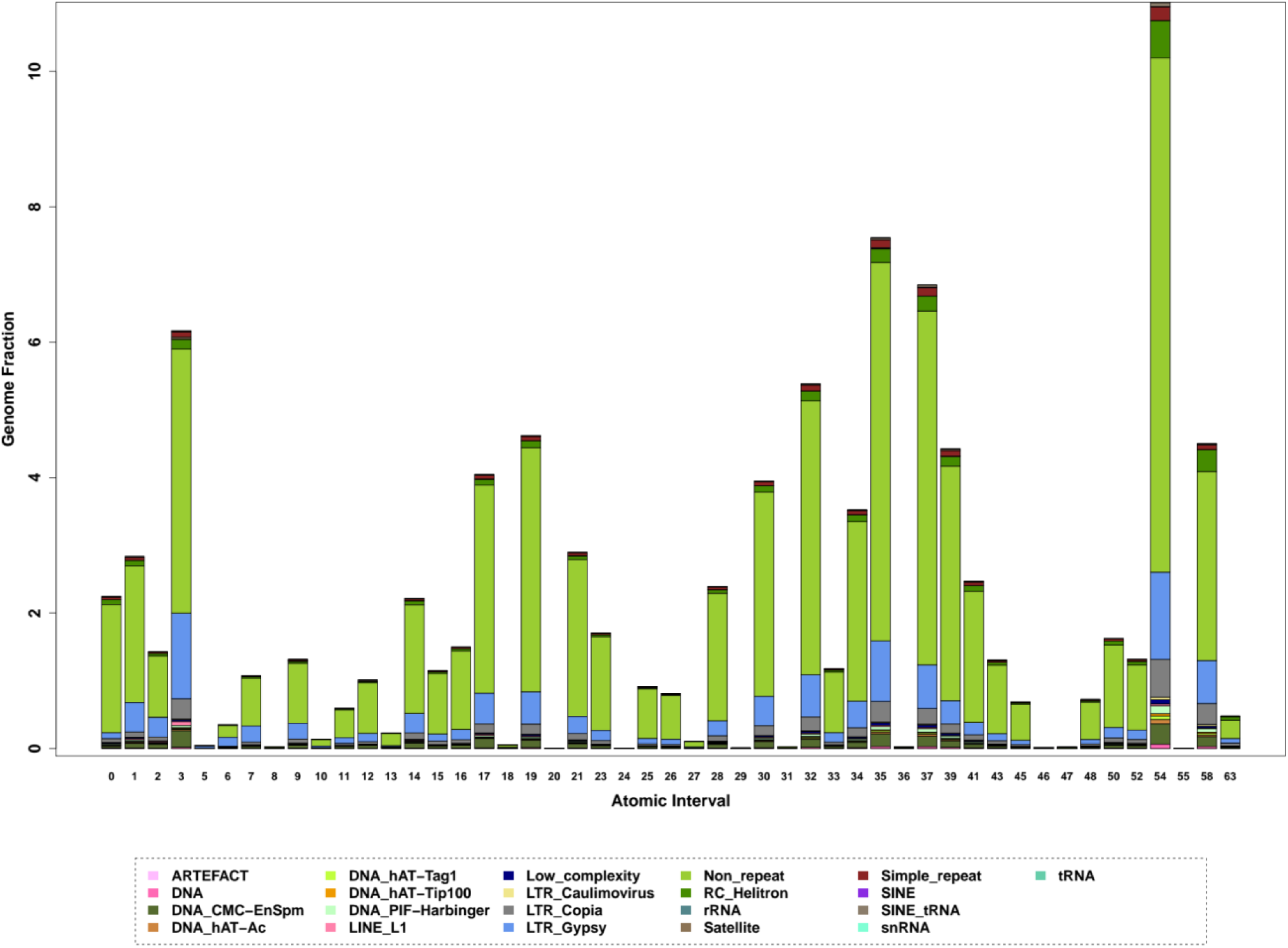
Fraction of genome contributed to each atomic interval of *Populus trichocarpa* PSMC by the repeat classes. Contribution of various repeat classes (percentage of whole genomic length) to each atomic interval is shown. LTR-Gypsy has contributed to around 2% in the atomic intervals spanning recent and ancient times, which might be one of the contributing factors to the change in Ne trajectory.

Genomic regions are assigned to a specific atomic interval based on the TMRCA of that region. This leads to a trend of increasing levels of heterozygosity from atomic intervals that correspond to recent to older time points. Each of the repeat classes independently shows this trend of increase in heterozygosity similar to non-repeat regions (see **Fig. 3**). Comparable estimates of heterozygosity between repeat and non-repeat regions suggest that the heterozygous sites identified in repeat regions are not merely variant calling artefacts. The ratio (Ts/Tv) of the number of transitions (Ts) to the number of transversions (Tv) has been used to evaluate the accuracy of variant call sets (Wang et al. 2015). Regions of the genome with artefactual variant calls would have a Ts/Tv ratio very different from the genomic average. Hence, as an additional validation of the variants identified within repeat regions, we calculated the Ts/Tv ratio for each repeat class by the atomic interval. While the estimates of heterozygosity showed an increasing trend towards older atomic intervals, we found that the Ts/Tv ratio did not show any discernible trend (see **Fig. 3**). Similar estimates of the Ts/Tv ratio in repeat and non-repeat regions suggests that the heterozygous sites identified in repeat regions are truly polymorphic.

**Figure 3:**
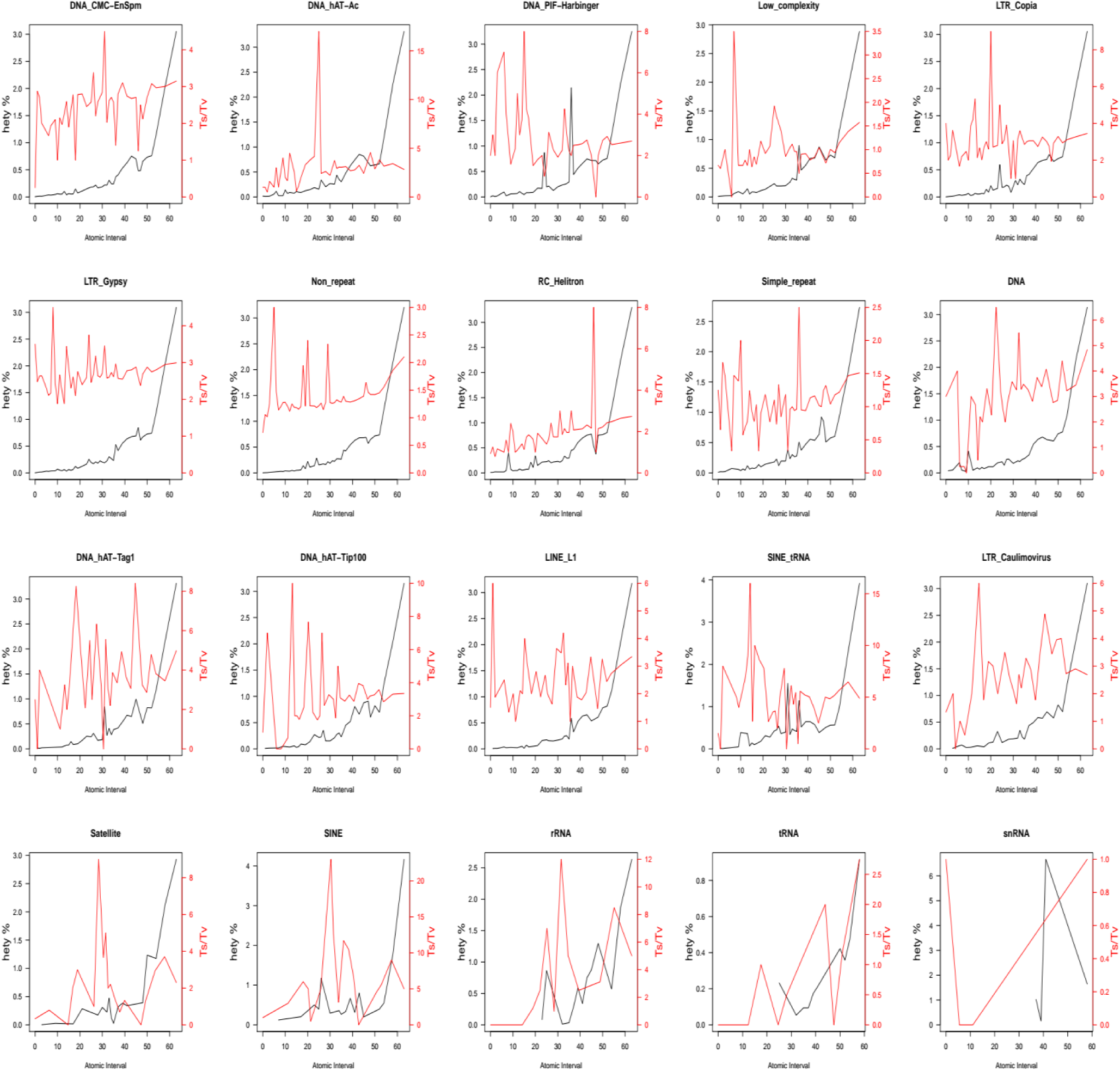
Comparison of heterozygosity and T_s_/T_v_ ratio across atomic intervals of Populus trichocarpa *PSMC*. Change in heterozygosity and corresponding T_s_/T_v_ ratio across atomic intervals in different repeat classes of *Populus trichocarpa* PSMC. The heterozygosity (left y-axis) increases with atomic intervals (x-axis) whereas T_s_/T_v_ ratio (right y-axis) does not follow this trend for most of the repeat classes.

Repeat regions of the genome have very high coverage due to the mapping of reads from multiple copies and low callability due to a large number of mismatches across the reads mapped to the same genomic region. Hence, some studies tend to mask genomic regions based on criteria determined based on coverage or callability instead of the presence of repeats. We separately evaluated the effect of masking genomic regions based on coverage or callability classes (see **Fig. S3 and S4**) and find that masking based on these criteria needs to be treated independently of repeat region-based masking.

### Which biological and technical factors influence demographic inference?

Several technical factors such as the optimal PSMC parameter settings, sequencing platform used, the prevalence of cross-contamination from closely related species, misleading or incomplete metadata in public datasets are important considerations during the comparative interpretation of demographic history. Similarly, biological factors such as the prevalence of whole-genome duplications, changes in the karyotype or genome size and high intraspecific variation in genetic diversity also need to be considered. We generated whole-genome sequencing data for a tropical plant species (*Mesua ferrea*) and use it as an example to understand how biological and technical factors influence demographic inference. For any newly sequenced genome, the PSMC parameters –r (initial theta/rho ratio), -p (pattern of parameters specifying distribution of free intervals and atomic intervals) and –t (the maximum time to TMRCA) need to be optimised so that all the atomic intervals have sufficient number of recombination events.

### The maximum time to TMRCA

For the first PSMC run of the *Mesua ferrea* genome, -t 5 -r 5 -p “4+25*2+4+6” options were used. However, the resultant PSMC output did not have enough recombination events in some of the atomic intervals. Therefore, the -p parameter was optimized until all the atomic intervals had a sufficient number of recombination events. A mutation rate of 2.5e-09 per site per year i.e. 3.75e-08 per site per generation was used assuming 15 years of generation time for scaling the results. The resultant trajectory did not go back to older (i.e., beyond 150 Kya; see **Fig. 4a**) time points. So, we decided to optimize all the parameters so that the trajectory will give meaningful results beyond 150 Kya. Hence, the maximum time to TMRCA, i.e., -t parameter was increased so that the trajectory extended to older (i.e., 150 to 400 Kya; see **Fig. 4a**) time points. The -p parameter was optimized along with -t, as increasing -t gave less number of recombination events in some of the atomic intervals. While maintaining 64 atomic intervals, we were able to get a reliable demographic trajectory with options -t 65 -r 5-p “3+2*17+15*1+1*12” i.e. 64 atomic intervals distributed across 19 free intervals (1+2+15+1). To know how far the trajectory might be extended back in time if we increase – t, we used -t 500 for one run, which however did not have a sufficient number of recombination events in most of the atomic intervals.

To know how increasing maximum time to TMRCA was altering the Ne trajectory, we considered some of the longest scaffolds and visualised the assignment of specific genomic regions to various atomic intervals. Comparing the atomic intervals assigned to the same genomic region at different values of -t, we found that genomic regions which were assigned to older atomic intervals for smaller values of -t were assigned to relatively recent atomic intervals with an increase in the -t parameter (see **Fig. 4b**). This redistribution of regions with increasing values of –t can be better understood by looking at changes in the distribution of lengths of genomic regions assigned to each atomic interval (see **Fig. S5**). For instance, in the case of *Mesua ferrea* the length of older atomic intervals tends to decrease with increasing values of –t. Genomic regions contributing to the older atomic intervals at higher values of the –t parameter become shorter and highly heterozygous. We ensured that such short high heterozygosity regions are not merely variant calling artefacts by visualising the atomic intervals assigned to genomic regions along scaffolds with associated heterozygosity and callability at these regions (see **Fig. 4b**).

**Figure 4a:**
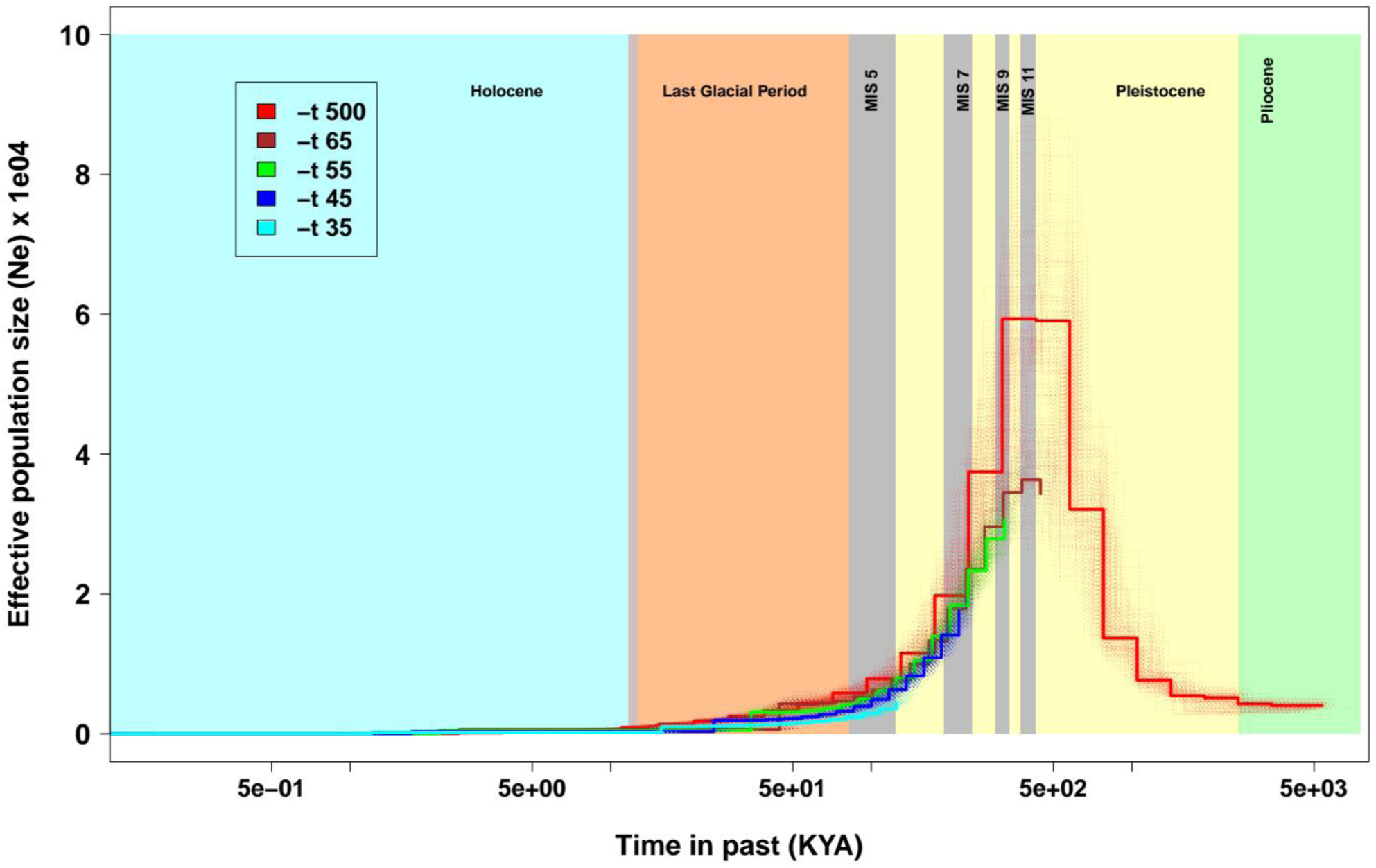
Demographic inference of *Mesua ferrea* by PSMC and effect of different values of maximum TMRCA. PSMC inferred trajectories with same -p parameter (3*2+1*10+15*2+14+4) but for several values of maximum TMRCA parameter. Colour used for -t of 35 (cyan), 45 (blue), 55 (green) and 65 (brown). For -t 500 (red), -p was used “4+25*2+4+6”, but did not have sufficient number of recombination events in some of the last atomic intervals. Demographic scenario shows steep decline in N_e_, after MPT (Mid-Pleistocene transition) i.e. ∼700 KYA, which again went through a second bottleneck during LGM (Last glacial maximum) of Last glacial period i.e. around ∼30KYA.

**Figure 4b:**
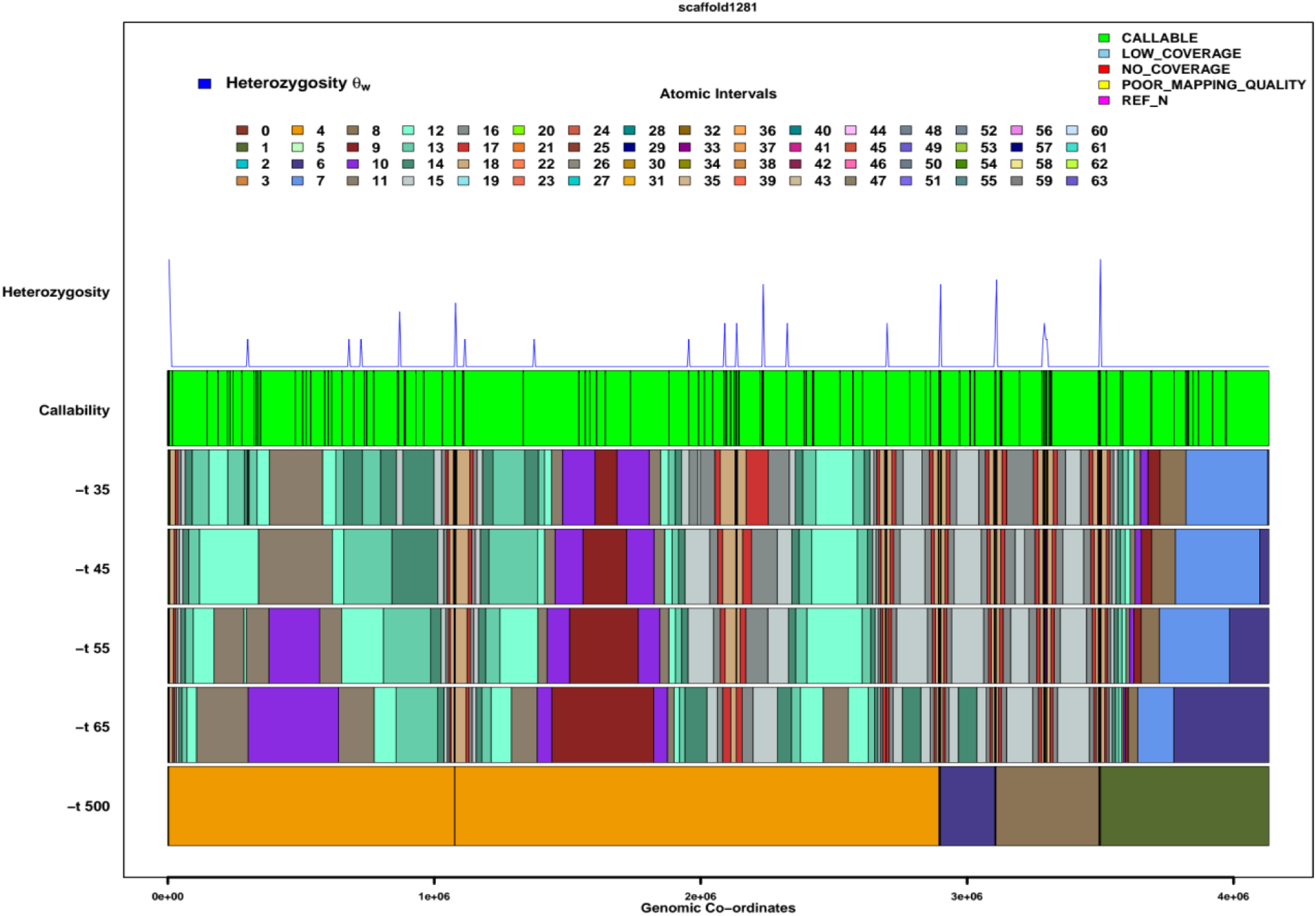
Distribution of atomic intervals across scaffold1281 for different maximum TMRCA values for *Mesua ferrea* PSMC. For each run of PSMC with different -t values decode based genomic regions along this scaffold and corresponding atomic intervals are shown. The atomic intervals which spanned scaffold1281 are shown here with their respective colours. Callability of bases in these regions is shown to highlight the quality of variants identified; heterozygosity is shown to demarcate hypervariable regions. It can be seen that same genomic coordinates are being distributed to more recent atomic intervals from older AI’s, which hints at redistribution of positions of atomic intervals with changes in the maximum TMRCA parameter values.

### The initial theta/rho ratio

The ratio of heterozygosity to recombination rate generally does not have an impact over the trajectory inferred using PSMC, but an increase in the value of -r does increases the number of recombination events and achieves convergence faster. We used the same -p and -t values (i.e., -t 65 -p “3+2*17+15*1+1*12”) with different values of –r. With decreasing values of –r, the Ne trajectory of *Mesua ferrea* extended further back in time (i.e., beyond 400KYA; see **Fig. S6a**). However, the atomic intervals corresponding to the extended trajectory did not have a sufficient number of recombination events for –r values less than 5. When the –r values were set at 5 or more, the convergence was achieved faster (see **Fig. S6b**).

### Does the sequencing platform affect the result?

We have previously shown that the trajectory of Ne can show extremely contrasting patterns between different populations of the same species (Vijay et al. 2018). To understand the variability in the demographic histories of different populations of *Mesua ferrea* we wanted to sample additional populations. Upon searching the European Nucleotide Archive (ENA), we found that a re-sequencing dataset labeled as *Mesua ferrea* sampled from Yunnan, China (see **Table S2**) was available for download. Surprisingly, the demographic trajectory inferred using this dataset extended back in time with -t of 5 (optimised with -r 5 -p “4+25*2+5*2”) and gave a different inference (see **Fig. S7**). However, we found that the sequencing for the sample from Yunnan had been performed using the BGISEQ-500 platform. In order to rule out the possibility of sequencing platform-specific technical issues, we compared the demographic trajectory of the Human individual NA12878 sequenced using BGISEQ-500 (see **Table S2** for dataset details) with the trajectory obtained for the same individual when the sequencing was performed using Illumina platform (see **Fig. S8**). We did not find any differences in the Ne trajectories estimated using BGISEQ-500 and the Illumina platform.

### Are the differences in Ne trajectories due to biological differences?

Having ruled out the possibility of sequencing platform-specific technical factors we considered the possibility of biological differences between the two samples of *Mesua ferrea*. Biological reasons for different trajectories can involve (a) different demographic histories of specific populations or (b) changes in the karyotype altering recombination landscape or (c) changes in genome size due to segmental or whole-genome duplication events. Since an earlier study has documented the prevalence of ecotype specific differences in the genome size of *Mesua ferrea* (Das et al. 2018), we first decided to compare the approximate in-silico genome size estimates of the two individuals under consideration. However, genome size estimates from raw read data are sensitive to the coverage depth, sequencing error rate, and polymorphism rate. Nonetheless, we see that the estimated genome size of the sample from China is (approximately 478 Mbp; see **Fig. S9 and S10**) ∼ 20Mbp less than the genome size estimated for the sample from India. We also observed that the heterozygosity of the sample from China was ∼25 fold higher than the sample from India. These differences in heterozygosity could be attributed to multiple factors such as contamination of reads from other species, independent Whole Genome Duplication (WGD) in Chinese sample, incorrect metadata regarding species identity in the public sequencing data repository. However, the WGD program (Zwaenepoel and Van De Peer 2019) used to detect whole-genome duplication events from genomic data did not find any evidence to support a WGD event specific to the Chinese sample based on the distribution of synonymous substitution rates (Ds) (see **Fig. S11**). Despite ruling out the possibility of bias from sequencing technology, we are not able to conclusively establish the reasons for the difference in the Ne trajectories due to reasons beyond the scope of this study.

### Comparative demography using PSMC on forest plant genomes

Estimation of the demographic history of multiple species of forest plants can provide useful information about the overall evolution of forests and the role of ecological processes or climatic events. We collated publicly available forest plant genomes that had sufficient data and compared the demographic histories inferred using PSMC. Demographic trajectories of the tropical species showed a considerable decline in Ne during 300 KYA – 1 MYA (see **Fig. 5**). This decline in Ne corresponds to a common event irrespective of the species-specific population dynamics. The period which shows bottleneck in all these species might be attributed to the environmental conditions of this period. During this period two major glaciations and extensive de-glaciation events have been recorded, which were considered to be longer and harsher than normal (van der Hammen 1974; Verbitsky et al. 2018). Glaciations following the Mid-Pleistocene transition (MPT) changed the duration of glacial events from 41 KYs to 100 KYs, translating into longer dry and colder conditions (Pisias and Moore 1981; Clark et al. 2006). These dry environments also affected precipitation in the tropics, leading to a decrease in CO2 concentration and less rainfall in these regions (Hewitt 2000; Dupont et al. 2001; Clark et al. 2006; Cabanne et al. 2016). In contrast, the silver birch (*Betula pendula)* showed an increase in Ne during this period, which could be explained by its adaptation to the dry, cold and high-altitude conditions (Salojärvi et al. 2017). Based on our analysis of 15 forest plants we could infer that late Pleistocene glaciations had a great impact on most of the forest plant species leading to reduced Ne during this period.

**Figure 5:**
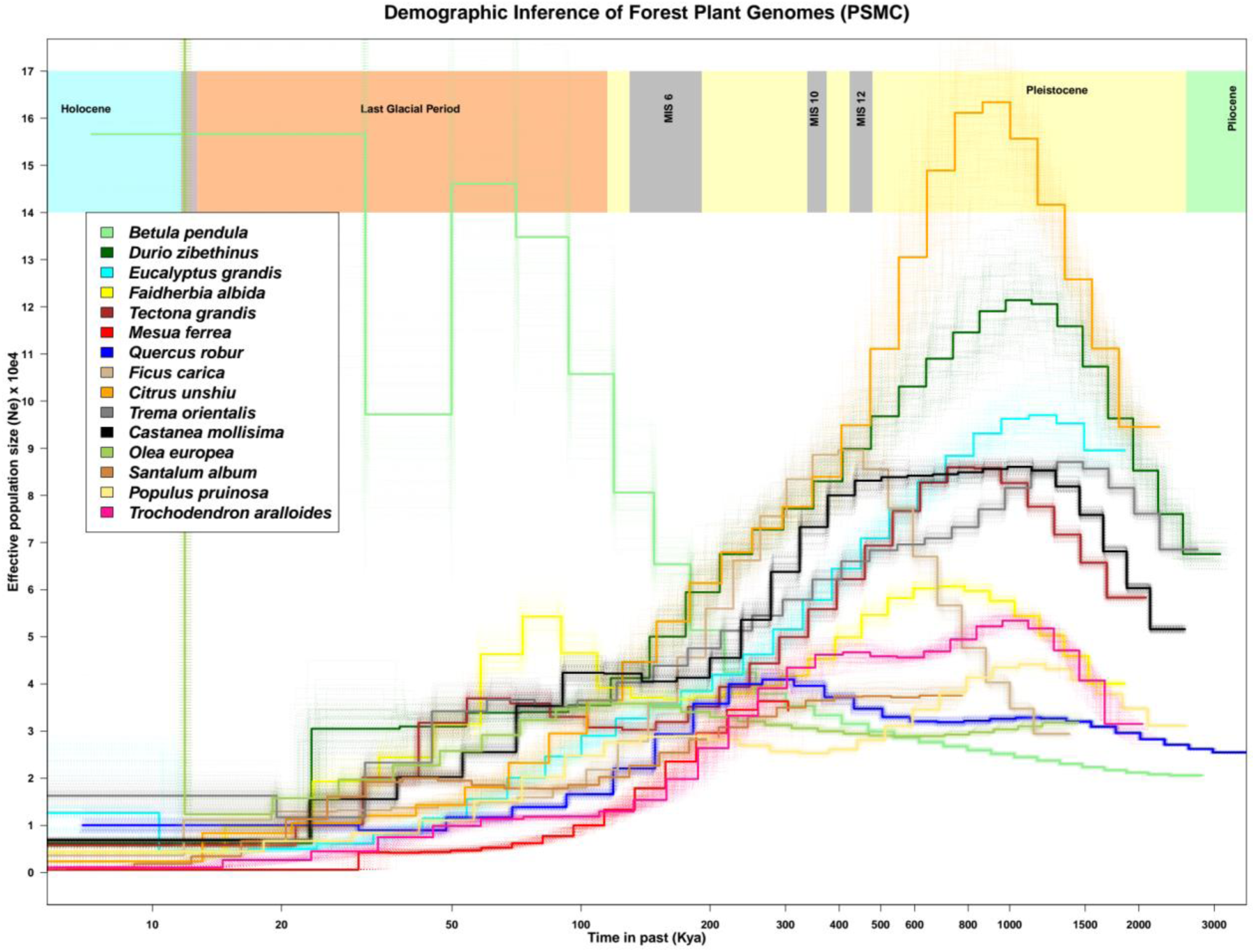
Comparative PSMC for Forest plant genomes. PSMC inferred trajectories with bootstrap replicates of 15 forest plant species. Top rectangles show respective time periods with important predicted glaciation events. *Betula pendula* shows a completely discordant trajectory compared to all other species. Whereas, tropically distributed species have a common trend of decline during and after Mid-Pleistocene glaciations. Some of the species such as *Faidherbia albida* were able to recover from these bottlenecks, which translates into their adaptation to dryer environments but most of the other plants were not able to recover from the same.

## Discussion

Using genomes and re-sequencing datasets of several species, we here explore how genome quality (contiguity, prevalence of gaps, assembly completeness), repeat abundance, technological heterogeneity (sequencing platform used, parameter settings optimised) and biological factors (changes in genome size) can affect the inference of demographic history. The scripts used for our analysis are implemented in the form of a re-usable pipeline with detailed documentation. We are thus confident that these quality control strategies and the associated pipeline will prove useful while comparing demographic trajectories between species to obtain insights into the underlying processes. The following paragraphs highlight our major findings and their relevance with respect to previous studies.

### Genome quality

We demonstrate that the estimates of Ne for the same sequencing dataset can be drastically different when the quality of the reference genome assembly changes. A recent study by Patton et al. (2019) investigated the robustness of several demographic inference methods to genome assembly quality and find that in comparison to other methods, PSMC robustly estimates Ne except in recent time periods. In contrast to our results, Patton et al. (2019) concluded that demographic inference methods are robust to the quality of the genome assembly. However, Patton et al. simulated differing amounts of genome fragmentation by manipulating the variant call file and overlook the possibility of genome quality-related biases introduced during the read mapping and variant calling steps. Moreover, these simulations assumed that random fragmentation of the genome would capture the complexity of differences in the qualities of real genomes. The ends of contigs or scaffolds in genomes are regions that are difficult to assemble, such as repeat-rich or hypervariable regions and are not randomly distributed in the genome. We consistently see differences in the demographic history estimated from different genome assembly versions of the same species. Yet, the demographic histories estimated from the 2012-devil and 2019-devil assemblies in Patton et al. are very similar. The negligible improvement (maximum difference in the percent of reads mapped is 0.01 considering 12 individuals) in the percentage of reads mapping to the recent version of the devil genome compared to the older version might explain why genome quality does not appear to influence the results in the case of the Tasmanian devil. Depending on the complexity of the genome architecture, the extent of improvement in genome assembly quality and which aspects of genome quality are improved will affect the magnitude of change in estimates of Ne. Hence, we urge caution while making generalisations regarding the effect of genome quality on the estimation of Ne.

### Repeat regions

Our results demonstrate that the inclusion of repeat regions does affect demographic inference but not all types of repeats have the same effect. The effect that a particular repeat class would have on the inference seems to depend on the abundance and genomic distribution of that particular repeat class. Hence, lineage-specific repeat classes can potentially affect the comparative analysis of demographic histories of closely related species. For instance, the LTR content might differ between closely related species (Zhang et al. 2020) and can heavily influence the results. We implement a quality control strategy that involves the comparison of demographic histories inferred from each repeat class separately. A better understanding of repeat class-specific mutation rates might allow for scaling each repeat type with an appropriate mutation rate and resolve this heterogeneity in the trajectories. In order to evaluate the effect of diverse repeat classes on the estimation of Ne, our pipeline relies on the existence of a reasonably good quality of repeat annotation in the focal species. While this is a caveat, it is a compromise done in order to finish the execution of the program in a timely manner. The users also have the choice to decide the number of repeat classes by combining repeat classes or separating them into sub-classes. A larger number of repeat classes leads to an increase in the runtime. We urge users to perform their own repeat annotation, identification and classification to overcome this limitation.

### PSMC parameter settings

Using the newly generated genome sequencing dataset of the tropical plant *Mesua Ferrea*, we demonstrate the effect of changing the three main parameter settings of the PSMC program. Our results highlight the importance of appropriately choosing the –t parameter (i.e., the maximum time to TMRCA) and provides intuitive understanding about changes in the distribution of genomic regions into specific atomic intervals as the values of –t is changed. The comparison of demographic histories across species requires that the PSMC parameter settings are properly optimised to identify relevant differences in their Ne trajectories. By comparing the trajectories of *Mesua Ferrea* individuals from two different populations, we show the importance of the –t parameter.

Certain historical time periods might be of greater importance due to specific climatic or geological events that have occurred in that time frame. Hence, it can be desirable to have a greater resolution while estimating the effective population size histories during these time periods. While increasing the number of free intervals in a particular time period will increase the resolution it can also lead to certain atomic intervals having too few recombination events. Hence, the atomic intervals need to be distributed such that each atomic interval has more than 10 recombination events after the 20th iteration. We hope that our results provide some clarity regarding the strategies to be used while choosing the –p parameter.

The output of PSMC is insightful when it is scaled to time in years based on the mutation rate and generation time of the species under consideration. While changes in the scaling parameters have been shown to result in similarly shaped trajectories it does change the absolute values of the estimates (Nadachowska-Brzyska et al. 2015). However, accurate estimates of mutation rates are missing or unreliable in the case of many species. Moreover, the estimates of mutation rates obtained by different methods can produce drastically different values. Generation time can also be difficult to estimate especially for long-living plant species. Hence, recent studies have resorted to scaling the results using multiple combinations of mutation rates and generation times to ensure the robustness of their observations.

## Conclusion

In summary, our study systematically investigates multiple sources of bias that can affect the inference of demographic history from whole genomic datasets. By comparing the demographic inferences obtained using different versions of the human, red flour beetle and zebrafish genomes, we establish that genome quality does have a considerable impact on the estimation of effective population size (Ne). Instead of simply masking repeat regions of the genome, we investigate the consequences of including each repeat class using the genome of the plant *Populus trichocarpa*. Interestingly, we find that most repeat classes are able to provide inferences consistent with those obtained from non-repeat regions and can be a viable source of demographic history. Our analysis of repeat regions is of special relevance as the quality of genome assemblies continues to improve with long-read sequencing technologies being able to correctly assemble repeat regions. Demographic histories inferred for the same human individual using BGI-seq and Illumina datasets show highly concordant results. Future studies that plan to compare datasets originating from these technologies are likely to benefit from our comparative analysis. Parameter settings used for running PSMC have also been investigated in considerable detail with guidelines for optimal use of parameters. Having investigated various sources of technical variability and parameter settings, we consider the genomic datasets of 15 forest trees and compare their demographic histories as a case study. We not only provide guidelines for performing filtering of genomic datasets but also develop the CoalQC pipeline that we hope will become a standard part of quality control prior to demographic analysis using whole-genome datasets.

### Material and Methods

#### Genome quality assessment

##### Assembly and annotation

Leaves of the plant *Mesua ferrea* were collected from a tree located near the College of Forestry, Ponnampet (GPS coordinates 12°08’56.5“N75°54’32.5”E; Altitude: 829-850 meter Above Sea level). Genomic DNA was extracted from the leaves using the standard CTAB protocol. The quality and integrity of DNA were assessed using 1x gel electrophoresis, Nanodrop, and Qubit. Illumina short-read (150bp) libraries were prepared with an insert size of 450 ± 50 bp to generate ∼110 Gbp of paired-end short-read data. The quality of the sequencing read data was assessed using the program FASTQC (Andrews et al. 2015). Genome size was estimated using Jellyfish (Marçais and Kingsford 2011) followed by GenomeScope (Vurture et al. 2017) using k-mer size of 21. GenomeScope estimated genome size of approximately 496.72 Mbp. Despite the observation of variability in genome size between ecotypes of *Mesua ferrea*, flow cytometry-based estimates have consistently predicted genome size of approximately 684.5 Mbp (Das et al. 2018).

For assembling the genome, MaSuRCA (Maryland Super Read Cabog Assembler) (Zimin et al. 2017) was used with parameters USE_LINKING_MATES = 1 CLOSE_GAPS=1 NUM_THREADS = 32 JF_SIZE = 12000000000 SOAP_ASSEMBLY=1. Our genome assembly length of 614.35 MB is slightly less than but comparable to previous estimates based on flow cytometry. To assess the quality of the assembled genome, we employed multiple quality assessment methods. While N50, N75, etc. are accepted metrics of genome contiguity, the number of genes assembled in the assembly serves as a metric of assembly completeness. The amount of repeats assembled is also one of the relevant metrics which informs about the quality of the assembly in low-complexity regions. The Quast (Mikheenko et al. 2018) program was used to calculate assembly statistics i.e. N50, N75, Number of Ńs per 100 KB, etc. (see **Table S3**). BUSCO (Benchmarking Universal Single-Copy Orthologues) (Simão et al. 2015) was used to assess the completeness of the assembly, using eudicotyledons_odb10 (see **Table S4 and S12**) and embryophyta_odb10 (see **Table S5**) dataset together with previously sequenced genomes of order Malpighiales. LTR_retriever’s LAI module was used to determine assembly quality based on the LAI (LTR Assembly Index) score which assesses repeat content assembled (see **Table S6**).

Annotation was carried out using MAKER-P (Campbell et al. 2014) version 2 with MPI. Repeat libraries obtained from RepeatModeler (Smit, AFA, Hubley 2015) and LTR-retriever (Ou and Jiang 2018) were concatenated and used to mask repeat regions of the genome. Published CDS dataset of *Populus trichocarpa* and concatenated multi-fasta of all available Malpighiales proteins were used as homology evidence for the first round of de-novo annotation. The results of the first round of annotation were then used for training SNAP (Korf 2004) and AUGUSTUS (Stanke et al. 2008) implemented in BUSCO. These predictions were used for the second round of annotation in MAKER-P. Iterative rounds of annotation were carried out for 5 rounds until no further improvement was observed as assessed by the AED (Annotation Edit Distance) values (see **Table S7**).

The raw sequencing read data was used to separately assemble the chloroplast genome using the NOVOPlasty (Dierckxsens et al. 2017) program. The Maturase K gene sequence of *Mesua ferrea* was used as a seed sequence and *Garcinia mangostana* chloroplast genome was used as a reference. The assembler uses seed sequence to find reads that cover this sequence and starts overlapped sequence assembly. The assembled chloroplast genome had two sets of contigs. The orientation of the contigs was determined by dot-plot analyses (see **Fig. S13**) with *Garcinia mangostana* and other Malpighiales chloroplast genome sequences. The full length of the assembled chloroplast genome was 161.4 Kbp long. It was then annotated using GeSeq and visualised using OGDRAW (see **Fig. S14**) implemented in CHLOROBOX (Greiner et al. 2019). For assembling the mitochondrial genome matR gene sequence of *Mesua ferrea* was used as a seed. Assembled chloroplast sequence was used for comparison and WGS raw reads were used in NOVOPlasty. The total assembled mitochondrial sequence was 20084 bp long (see **Fig. S15**).

### Datasets used

To demonstrate the utility of our quality control pipeline and generality of our observations across diverse taxa we used species from different phyla such as nematodes, plants, vertebrates, etc. The published genome assemblies used as reference genomes were downloaded from NCBI/UCSC genome browser (details provided in **Table S8**). We searched the European Nucleotide Archive (ENA) for genomic sequencing datasets and downloaded those datasets that had >20X coverage (see **Table S2**). The raw read datasets were mapped to corresponding unmasked genomes using the short-read aligner BWA-MEM (Li 2013) with default settings.

### Genome assembly quality comparison

The latest version of the Human genome assembly, i.e., hg38, was downloaded from Ensembl, whereas previous assemblies i.e. hg19, hg15, hg10, and hg4 were downloaded from UCSC genome browser (see **Table S8**). The genome assembly statistics i.e. N50 statistics and Number of N’s per 100 Kb were calculated using Quast. SAMTOOLS flagstat module was used to get mapping percentages for each assembly using mapped alignments of each assembly. To get assembly completeness statistics, BUSCO was used with dataset Mammalia_odb9 on each of the assemblies. Genome assembly quality was also assessed for red flour beetle (*Tribolium castaneum*) and zebrafish (*Danio rerio*) genomes. These comparative assembly quality statistics are available in **Table S1**.

### Inference of demographic history using PSMC Parameter settings

Variant calling was performed using SAMTOOLS and BCFTOOLS with the depth parameters for vcf2fq command of vcfutils.pl decided based on the mean coverage of the reads. The resultant fastq file of heterozygous sites was converted into psmcfa format using the fq2psmcfa program for bin sizes of 20, 50, and 100. For the first run of PSMC, options were set as -t 5, -r 5, -p “4+25*2+4+6”. The output was evaluated to see if a sufficient number of recombination events had occurred in each atomic interval. If there were some atomic intervals that did not have at least ten recombination events after the 20th iteration, then the -p parameters were modified. For example, if -p parameter “4+25*2+4+6” is set, it has 64 atomic intervals distributed across 28 free intervals i.e. (1+25+ 1+ 1). Changing the distribution of atomic intervals across free intervals, e.g., “8+25*2+2+4” would be the first step to see if enough recombination events are obtained. If not, then changing the free intervals, i.e., “**3**\*2+**1**\*10+**15**\*2+**1**\*14+**1**\*4” (written as **free intervals** * atomic intervals) was done to get sufficient recombination events. Only after obtaining a sufficient number of recombination events, the -p parameter was finalised. PSMC was run with the -d (decode) option for identifying genomic regions that contributed to each atomic interval. After obtaining the PSMC output, an appropriate mutation rate and generation time were used to generate the scaled plot using the psmc_plot.pl script. For bootstrapping analyses, the psmcfa file was first split into equal lengths of 5 MB, and was used for 100 runs of PSMC.

### Effect of repeat regions

The unmasked genomes were analysed to identify and annotate repetitive regions. For genome-wide identification of LTR’s, the program LTR-retriever was run using repeat libraries made by concatenating LTR harvest (Ellinghaus et al. 2008) and LTR_finder v 1.0.6 (Xu and Wang 2007) output. The RepeatModeler program was used for the de-novo identification of repeats. Both genome-wide LTR-retriever and RepeatModeler repeat libraries were concatenated and given as input to the RepeatMasker program. The tabulated output file of RepeatMasker was converted to bed format and used for further analyses.

Separate runs of variant calling were carried out using the unmasked and masked genomes followed by PSMC analyses. PSMC was run with -d option for both unmasked and masked datasets, and outputs were produced for three bin sizes (specified using the –s flag), i.e., 20, 50, and 100. For each run of PSMC, the decode2bed.pl script was used to obtain details of the atomic interval assigned to specific genomic regions. The prevalence of repeat regions in each atomic interval was assessed by intersecting the positions of the repeats with the positions of atomic intervals using BEDTools (Quinlan and Hall 2010). The genomic coordinates of heterozygous sites and ratio of transitions (Ts) to transversions (Tv) were obtained using the hetlist command of seqtk (Li 2015). Subsequently, repeat class-specific heterozygosity and Ts/Tv ratio in each atomic interval was calculated using the positions of repeat regions, decoded atomic intervals and genome-wide list of heterozygote sites as arguments to BEDTools. To evaluate the effect each individual repeat class would have on PSMC, one repeat class was unmasked at a time keeping all the other repeat classes masked prior to PSMC analyses.

### Effect of coverage and callability

Genome-wide per base depths were calculated using SAMTOOLS depth command. The read-depth information was used to assign cumulative coverage classes, i.e., bases having >0 read depth, >10 read depth, >20 read depth and so on. The genome could thus be divided into regions that correspond to specific cumulative coverage classes. GATK version 3.8.1 (McKenna et al. 2010) was used to get callability information using CallableLoci command. The callable status obtained from CallableLoci was used to divide the genome into regions of a specific callability. The effect these coverage based and callability based classes would have on PSMC inference was assessed by performing PSMC analyses separately on each of the classes after masking all genomic regions that were outside the class under consideration.

### Comparative PSMC of forest plant genomes

Published plant genome assemblies till November 2019 and their details were obtained from the plant genome database (available at https://www.plabipd.de/timeline_view.ep). From this list of published plant genomes, forest plants (i.e. excluding annual plants) species with >20X coverage were selected. The genomes and corresponding short-read data were downloaded from public repositories (see **Table S2 and S8** for details). PSMC analysis was performed on each of these species with default parameters i.e. –t5 –r5 –p “4+25*2+4+6”. A mutation rate estimate of 2.5e-09 per site per year which has been used for *Populus trichocarpa* in an earlier study (Bai et al. 2018) was used for all the species. Generation time for each species was obtained through a literature search. For each species per generation, mutation rates were obtained using corresponding generation times (**Table S9**).

## Acknowledgments

We thank the Ministry of Human Resource Development for fellowship to ABP, the Council of Scientific & Industrial Research for fellowship to SSS. NV has been awarded the Innovative Young Biotechnologist Award 2018 from the Department of Biotechnology and Early Career Research Award from the Department of Science and Technology (both Government of India). The computational analyses were performed on the Har Gobind Khorana Computational Biology cluster established and maintained by combining funds from IISER Bhopal under Grant # INST/BIO/2017/019, IYBA 2018 from DBT and ECRA from DST.

## Author contributions

NV conceived of the study based on inspiration from CGK and designed the study along with ABP & CGK. ABP conducted computational analysis and compiled the results with assistance from NV. SBN and RS collected the samples required for primary data generation. SSS extracted DNA from the leaf samples. ABP, NV, and CGK wrote the manuscript along with SBN, RS, and SSS. All authors approved the final draft.

## Supplementary figure legends

**Figure S1a:**
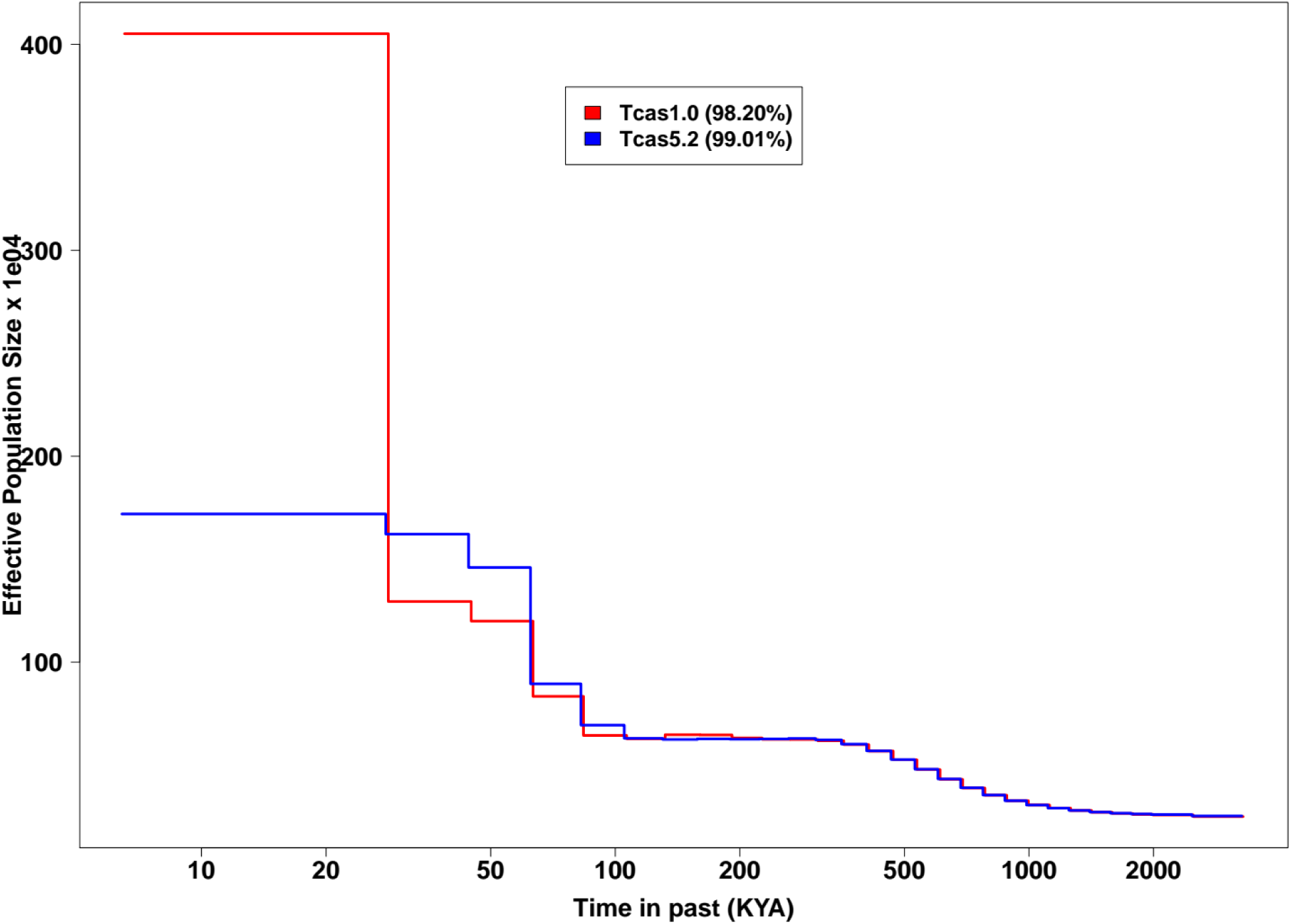
Change in Effective population sizes (N_e_) along with change in genome quality. Bootstrapped PSMC curves for *Tribolium castaneum* mapped to Tcas1.0 and Tcas5.2 genome assemblies with different genome quality. Mutation rate used 2.9e-09 per site per generation with generation time of 0.3 i.e. 12 weeks for one generation.

**Figure S1b:**
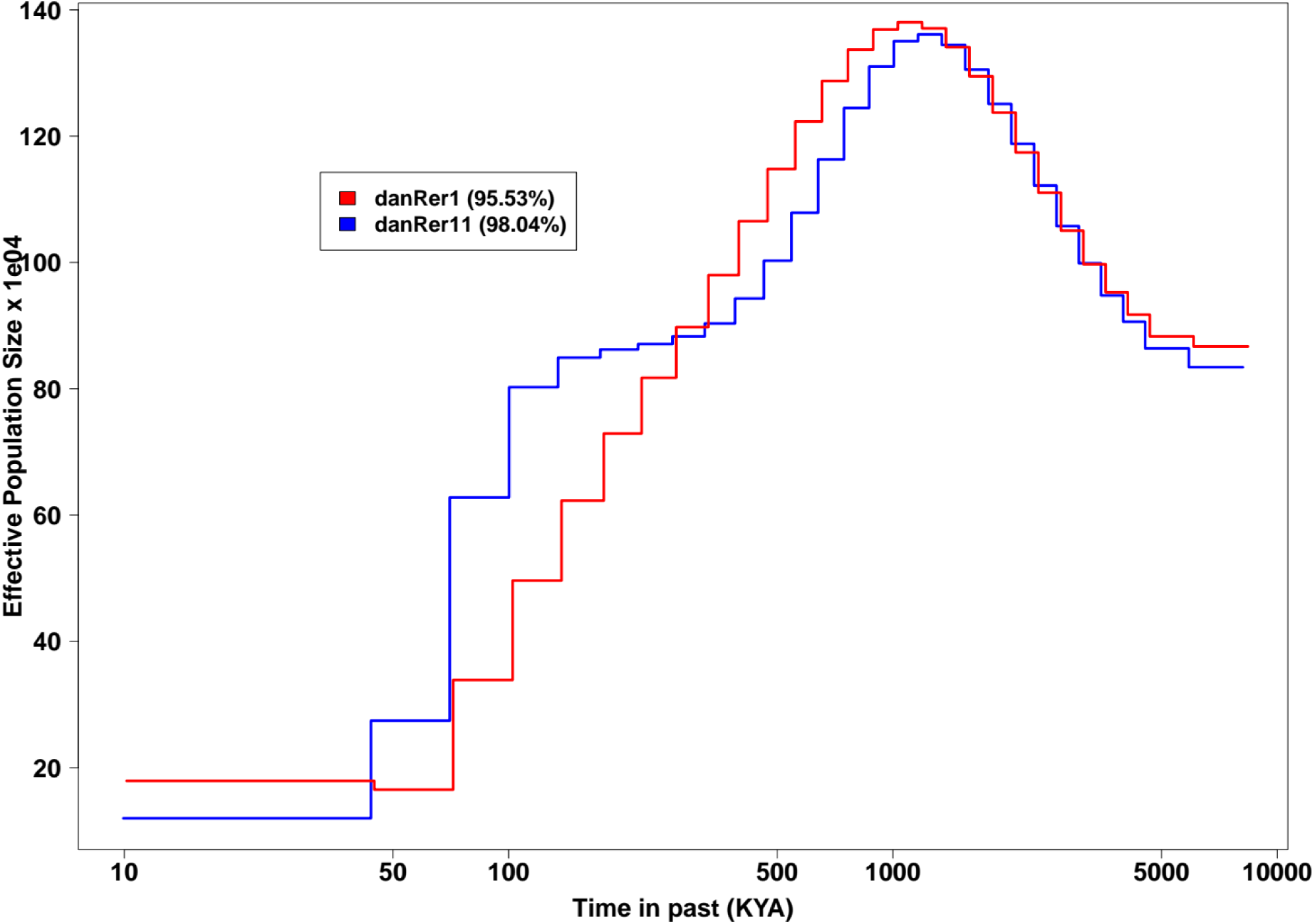
Change in Effective population sizes (N_e_) along with change in genome quality. Bootstrapped PSMC curves for *Danio rerio* mapped to danRer1 and danRer11 genome assemblies with different genome quality. Mutation rate used 1.9e-09 per site per generation with generation time of 1 year.

**Figure S2:**
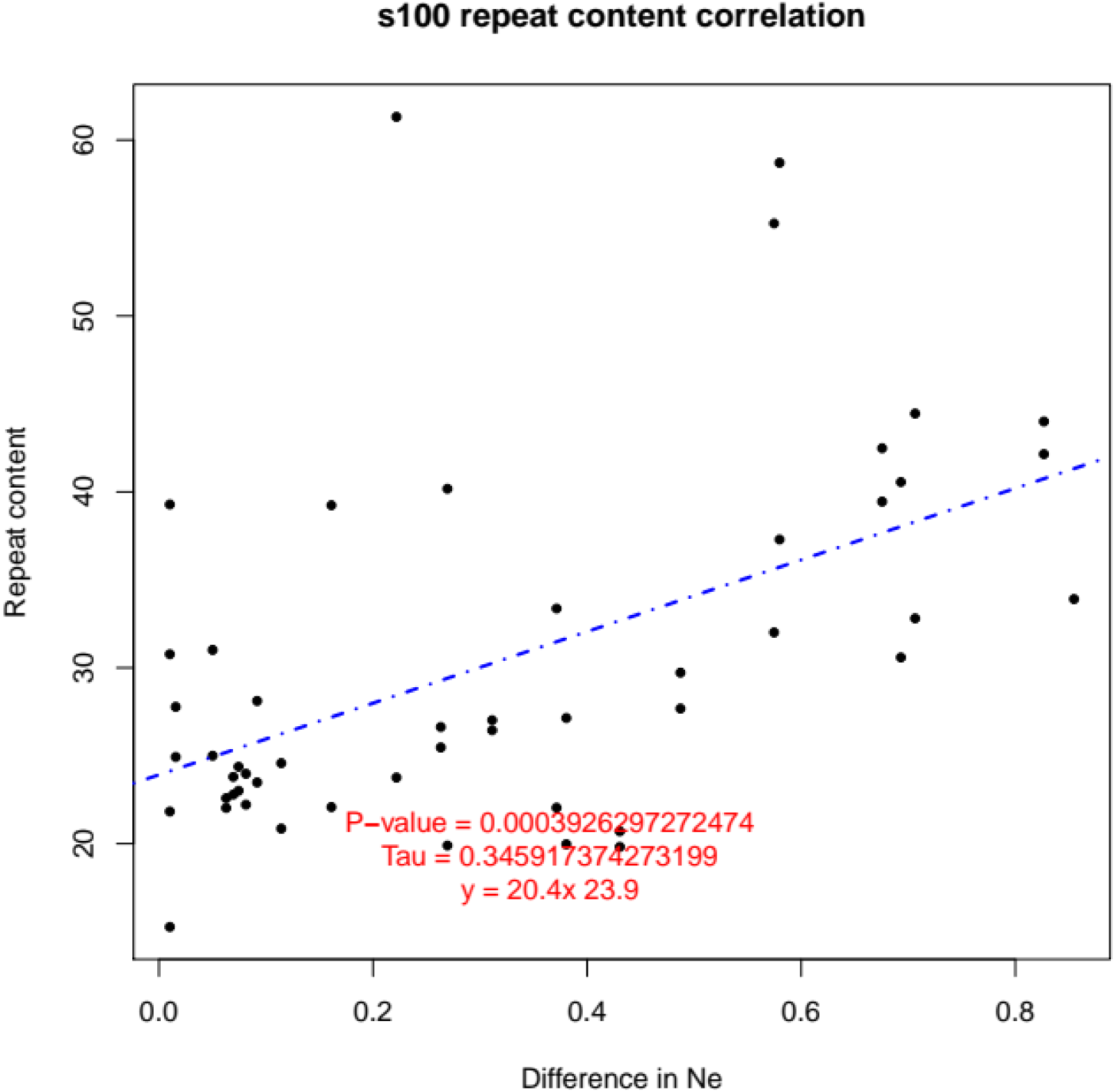
Correlation between repeat abundance and change in Effective population size (N_e_). Repeat content across atomic intervals in *Populus trichocarpa* PSMC showed a positive correlation with absolute change in N_e_ estimated from masked vs unmasked genomes. Kendall’s correlation coefficient was calculated showing Kendall’s correlation coefficient i.e. Tau (τ) =0.346, with p-value of 0.0004.

**Figure S3:**
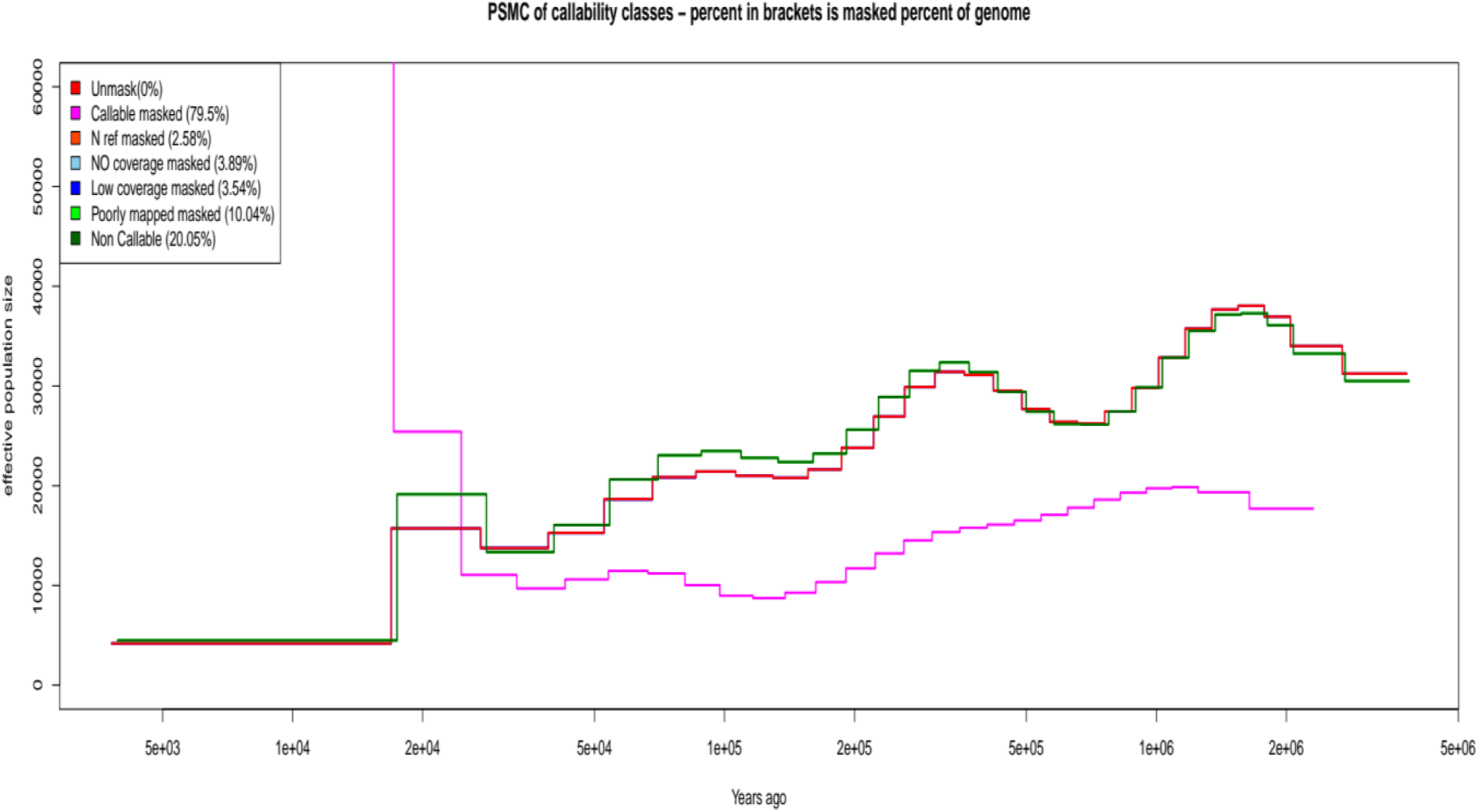
Effect of callability of bases on *Populus trichocarpa* PSMC. The output of CallableLoci module of GATKv3.8 distributed the genomic regions in several classes such as, Callable (good quality bases of reference genome), N-ref (Bases having N’s or gaps in the reference genome), No-coverage (Bases in the genome which were not supported by any read), Low-coverage (Bases in the genome which showed small support of reads compared to mean) and Poorly-mapped (Bases in the genome which showed poor mapping quality of reads). All of these classes were masked one at a time and the effect of each non-callable group was evaluated. After that all the non-callable groups were merged as a non-callable class and they were masked followed by another run of masking callable sites. For each run the percent of the genome masked is given in parentheses. Masking of callable sites gave completely different results, whereas other individual non-callable classes did not show any large change in the inferred trajectories. Non-callable trajectory showed some difference but did not change the trajectory much to change the inferences about the demography.

**Figure S4:**
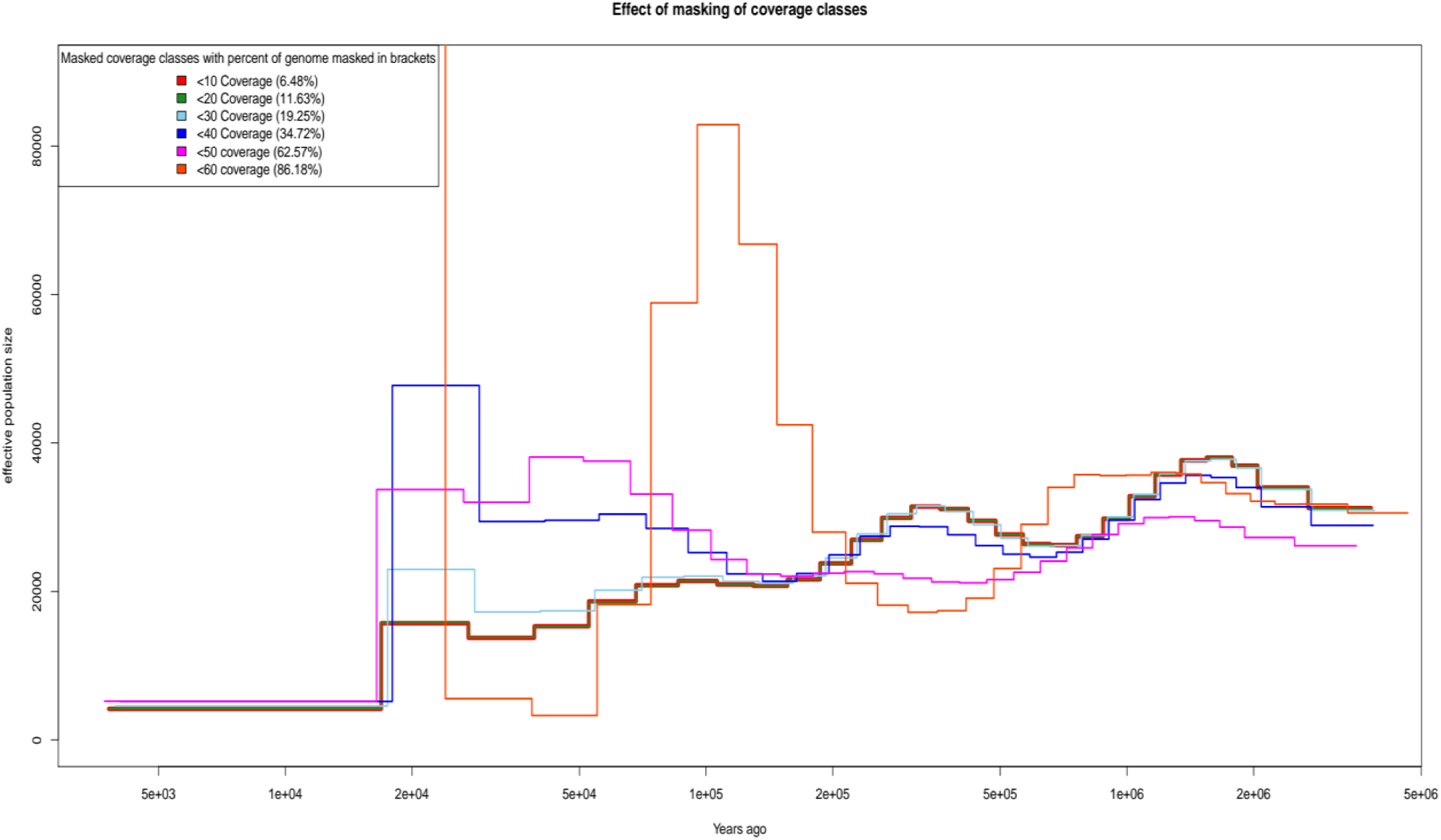
Effect of masking of different cumulative coverage classes on *Populus trichocarpa* PSMC. Whole genomic per base depth was calculated using SAMTOOLS depth command; this was used to make cumulative coverage classes based on their coverage distribution. Bases having < 10x coverage in one class, bases having < 20x coverage in another class etc. For each coverage class genomic co-ordinates were obtained and used for masking these regions, followed by PSMC analyses. The amount of genome masked by each class is showed in parentheses. There was no difference in PSMC trajectories till masking of less than 20X coverage classes, whereas from <30X coverage class masking trajectories started to differ till <60 X coverage class. Masking for more coverage classes i.e. < 70X and more did not produce psmcfa file during analyses. Genomic regions with higher coverages mostly contributed to the older atomic intervals, as masking till <40X coverage classes showed difference in recent time and small difference in older time.

**Figure S5:**
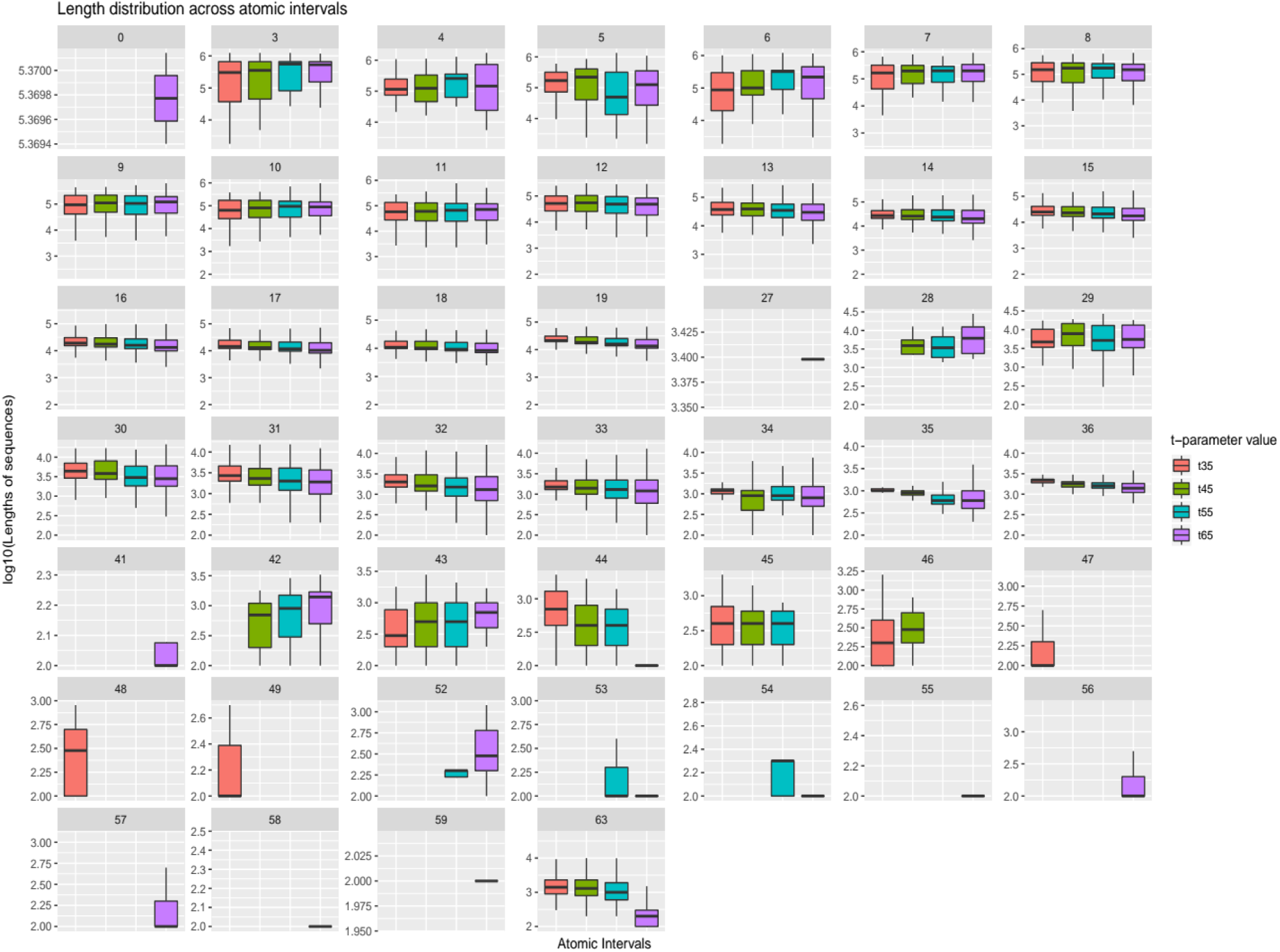
Change in length distribution of atomic intervals due to change in maximum TMRCA values in *Mesua ferrea* PSMC. Lengths of sequences in each atomic interval are compared for each maximum TMRCA value to evaluate if lengths are getting redistributed or not. For -t 65 (purple box) atomic intervals contributing to older times (see AI 53 to 63) are more represented than all other values, and even if present (see AI 53,54,55 and 63) -t 65 has smaller lengths compared to others in those time periods. This shows that increasing the maximum TMRCA allows shorter genomic regions to contribute to older times.

**Figure S6a:**
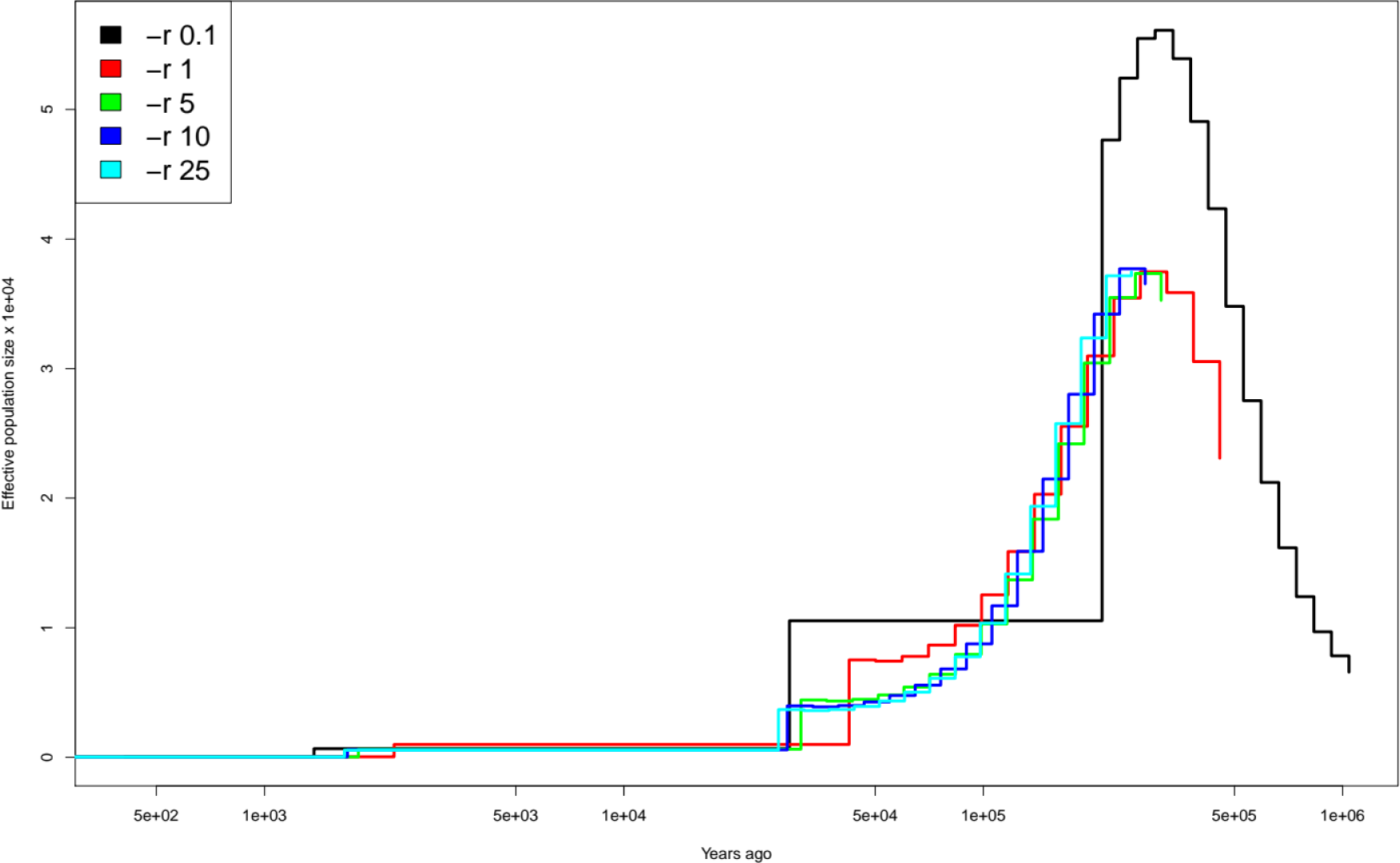
Effect of changing θ/ρ (r flag) value in PSMC on demographic inference of ***Mesua ferrea* PSMC.** PSMC estimates were inferred for *Mesua ferrea* with different values of -r flag. Smaller values of these value were able to travel further ahead in trajectory (see black and red lines), whereas atomic intervals contributing to these time points did not have enough recombination events. The other -r values i.e. 5,10 and 25 showed convergence in terms of recombination events but did not show any change in trajectory (see green, blue and cyan lines).

**Figure S6b:**
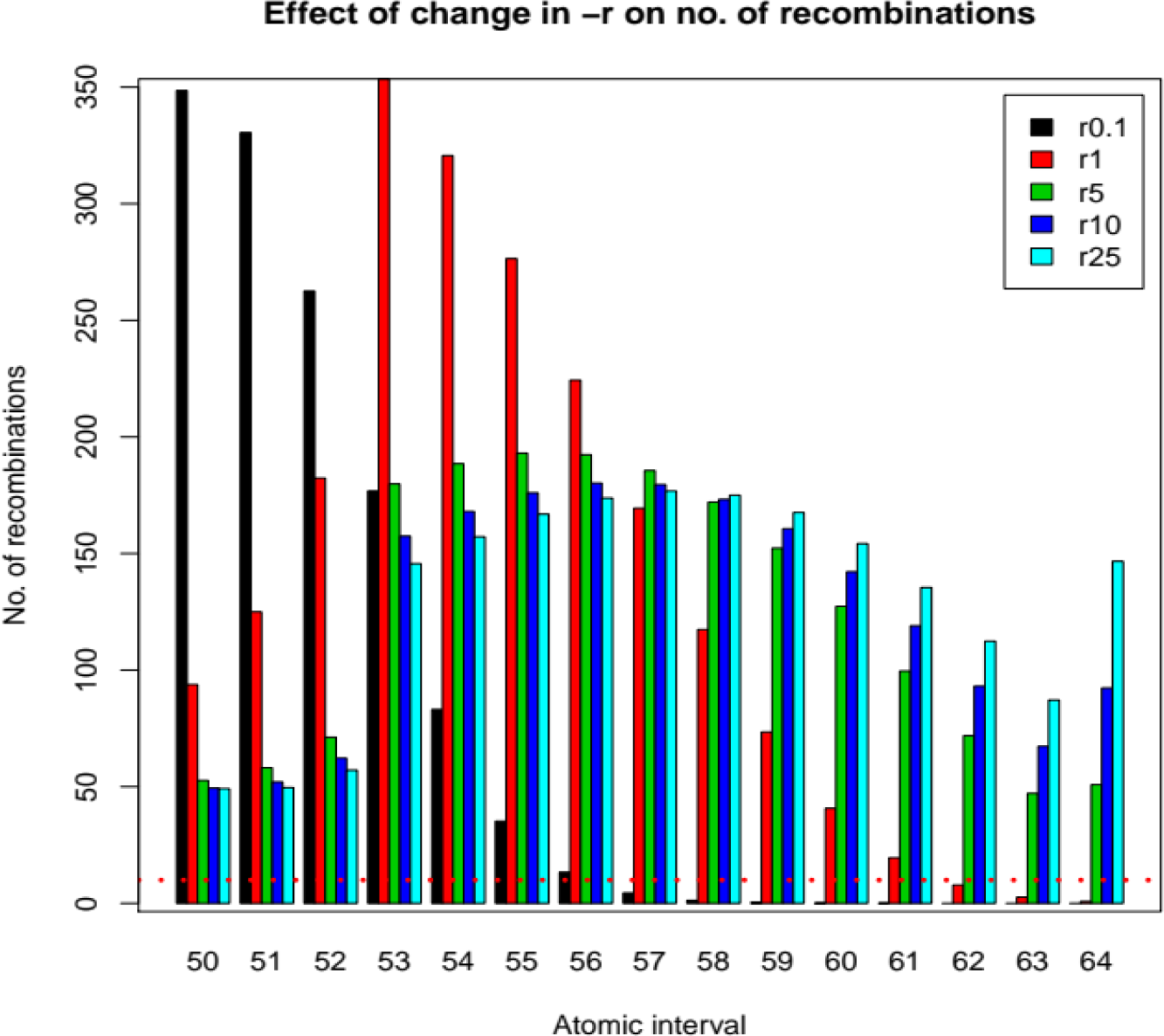
Effect of changing θ/ρ (r flag) value in PSMC on the number of recombination events across atomic intervals from 50 till 64 of *Mesua ferrea* PSMC. The -r values were able to go further ahead in trajectories with smaller values i.e. 0.1 and 1, but these atomic intervals did not show convergence in terms of recombination events. For the value of 0.1, till 56^th^ AI enough recombination events occurred, whereas for value of 1 there were less than 10 recombination events for last three atomic intervals.

**Figure S7:**
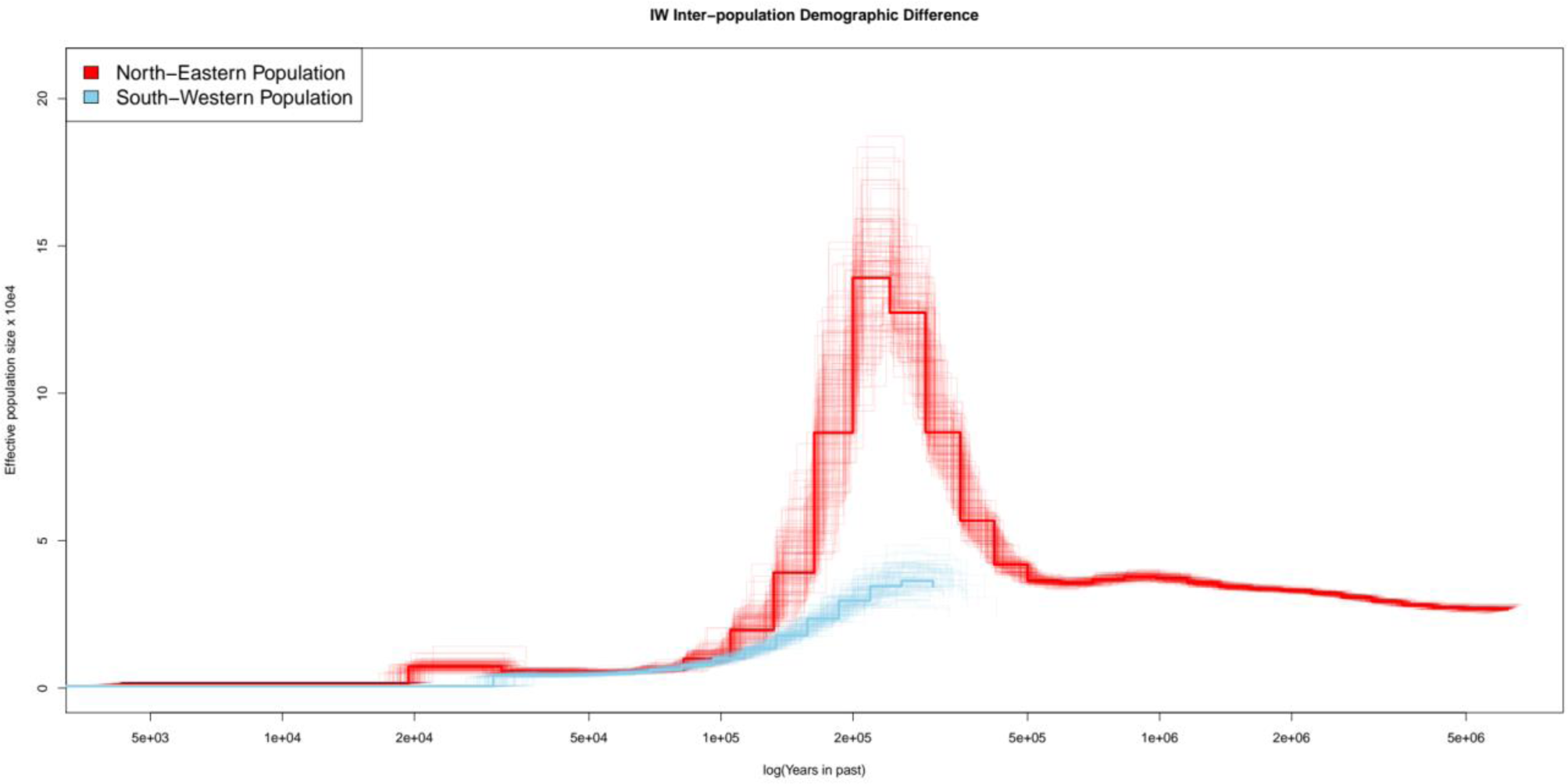
Demographic inference of *Mesua ferrea* for North-eastern (China) population (red) and South-western (India) population (sky-blue). Chinese sample (red) trajectory extends well back in time till ∼5MYA, whereas Indian sample (sky-blue) reaches till ∼400 KYA only. The time at which the population decline begins is similar in both and shows similar trajectories from ∼100 KYA till the recent times, as both show second decline during last glacial period i.e. ∼20 KYA.

**Figure S8:**
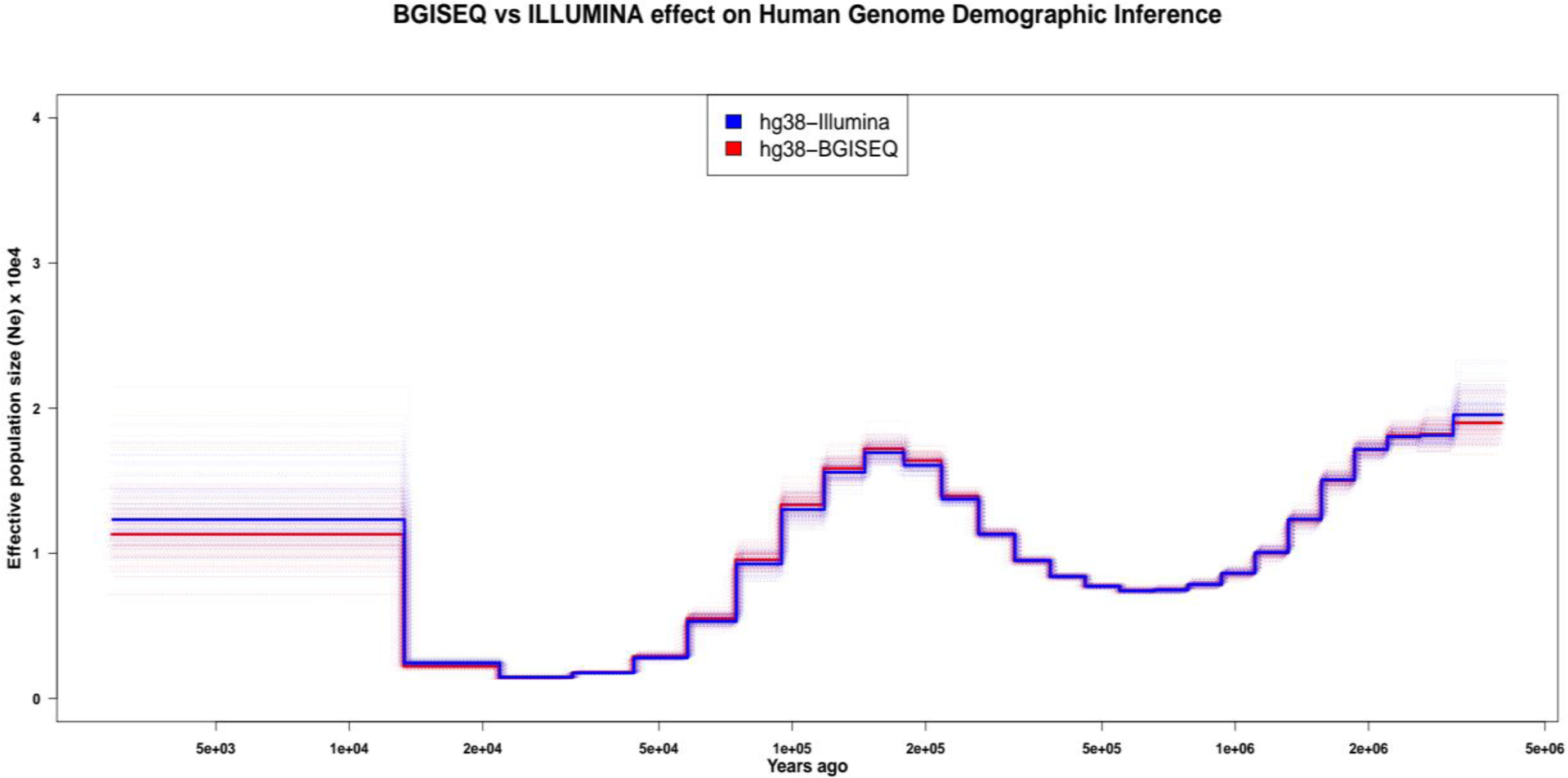
D**e**mographic **inference of Human-NA12878 sequenced using Illumina (blue) and BGISEQ (red).** The inferred PSMC trajectories showed identical Ne estimates, showing there are no sequencing platform based differences.

**Figure S9:**
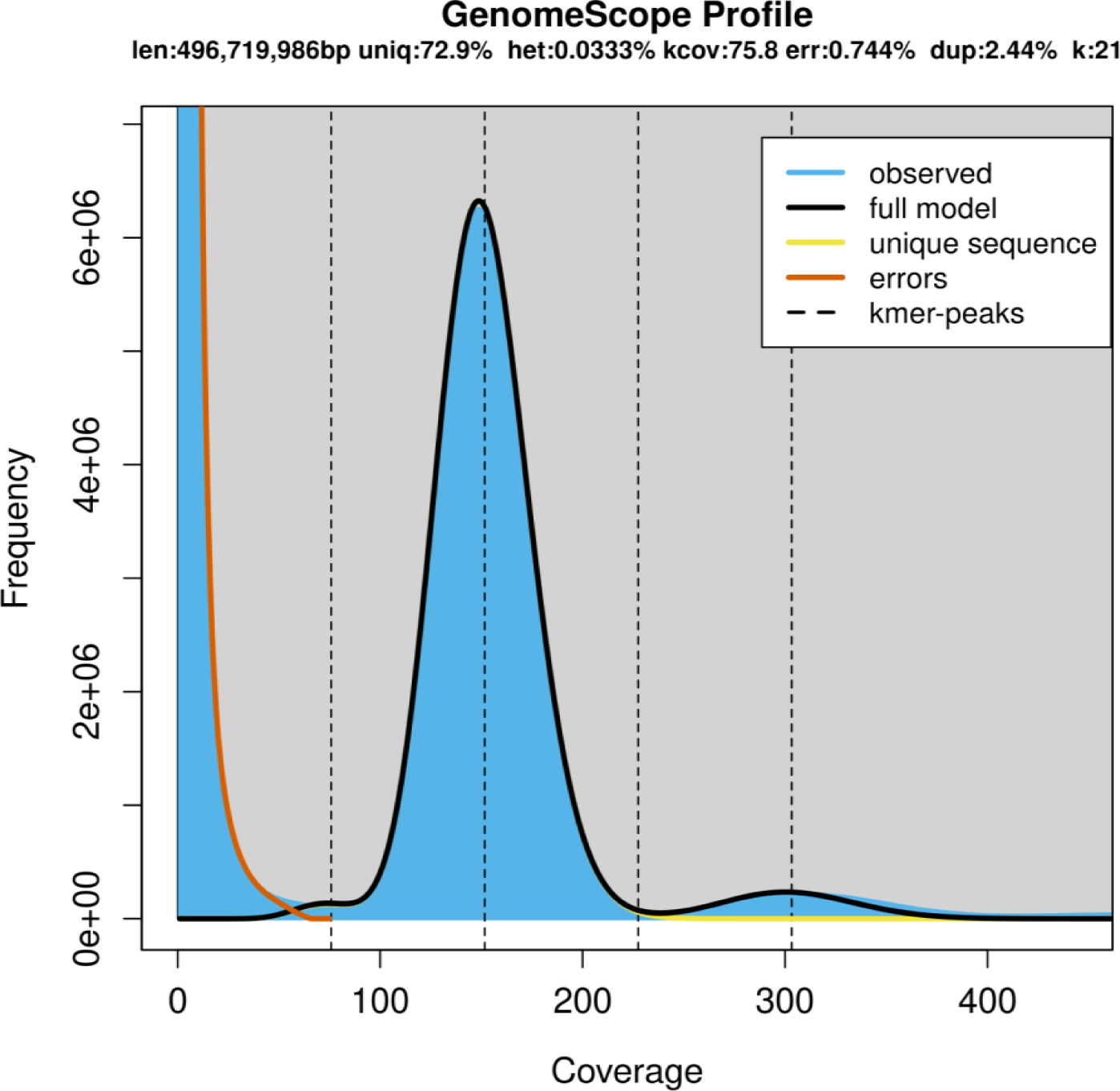
K-mer distribution of 21-mer’s of sample from Indian population *Mesua ferrea*. GenomeScope results of Indian sample predict low heterozygosity (0.03%) of the sample with a single peak at 150x coverage. Predicted genome size is approx. 497 Mbp, which is underestimation owing to neglecting high coverage sequences of organellar and repeat sequences.

**Figure S10:**
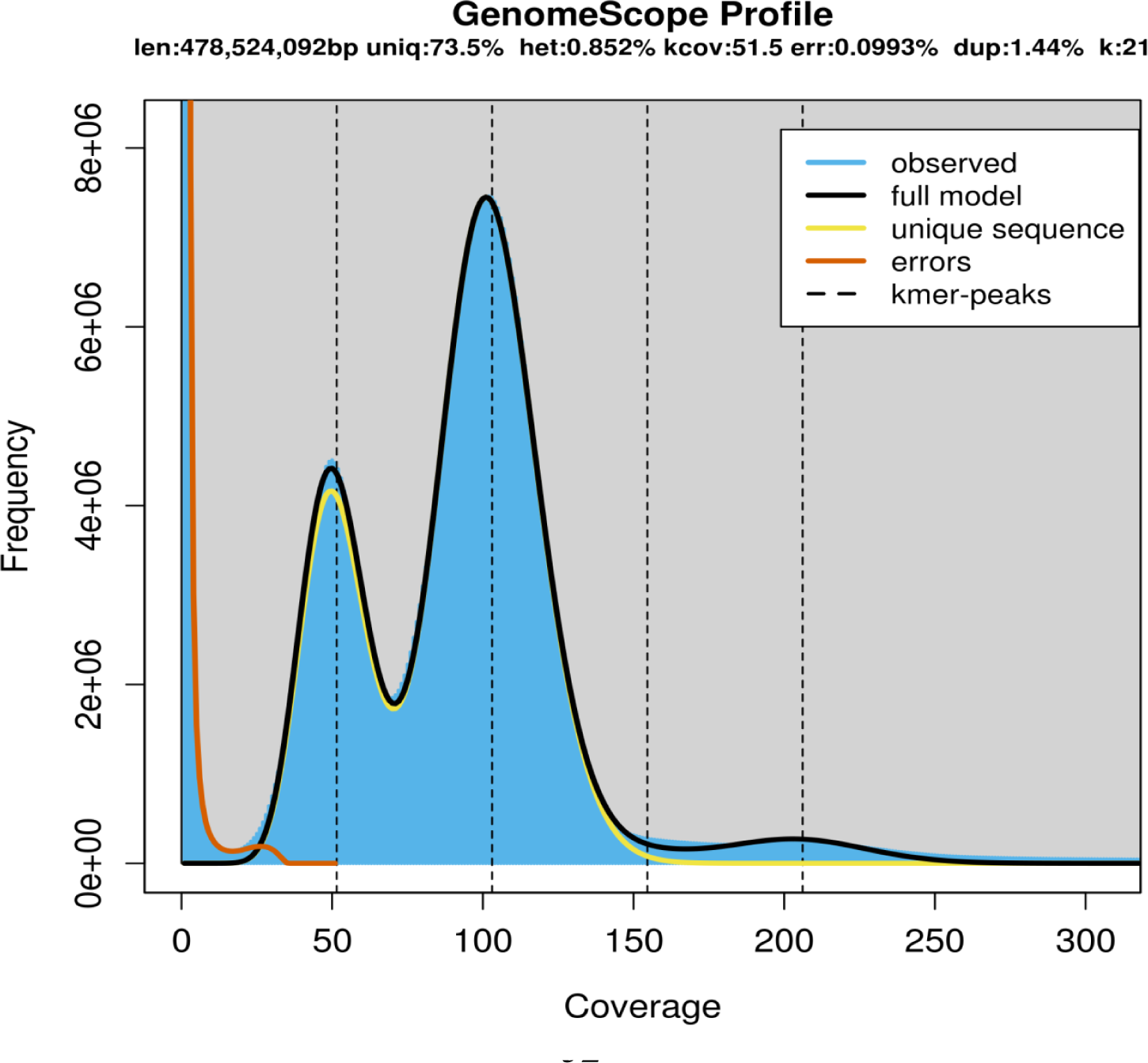
K-mer distribution of 21-mer’s of sample from Chinese population *Mesua ferrea*. GenomeScope results of Chinese sample predict high heterozygosity (0.85%) compared to Indian sample, showing ∼25-fold difference between both populations. Two peaks are due to high heterozygosity, but predict somewhat similar genome size i.e. 479 Mbp compared to other sample.

**Figure S11:**
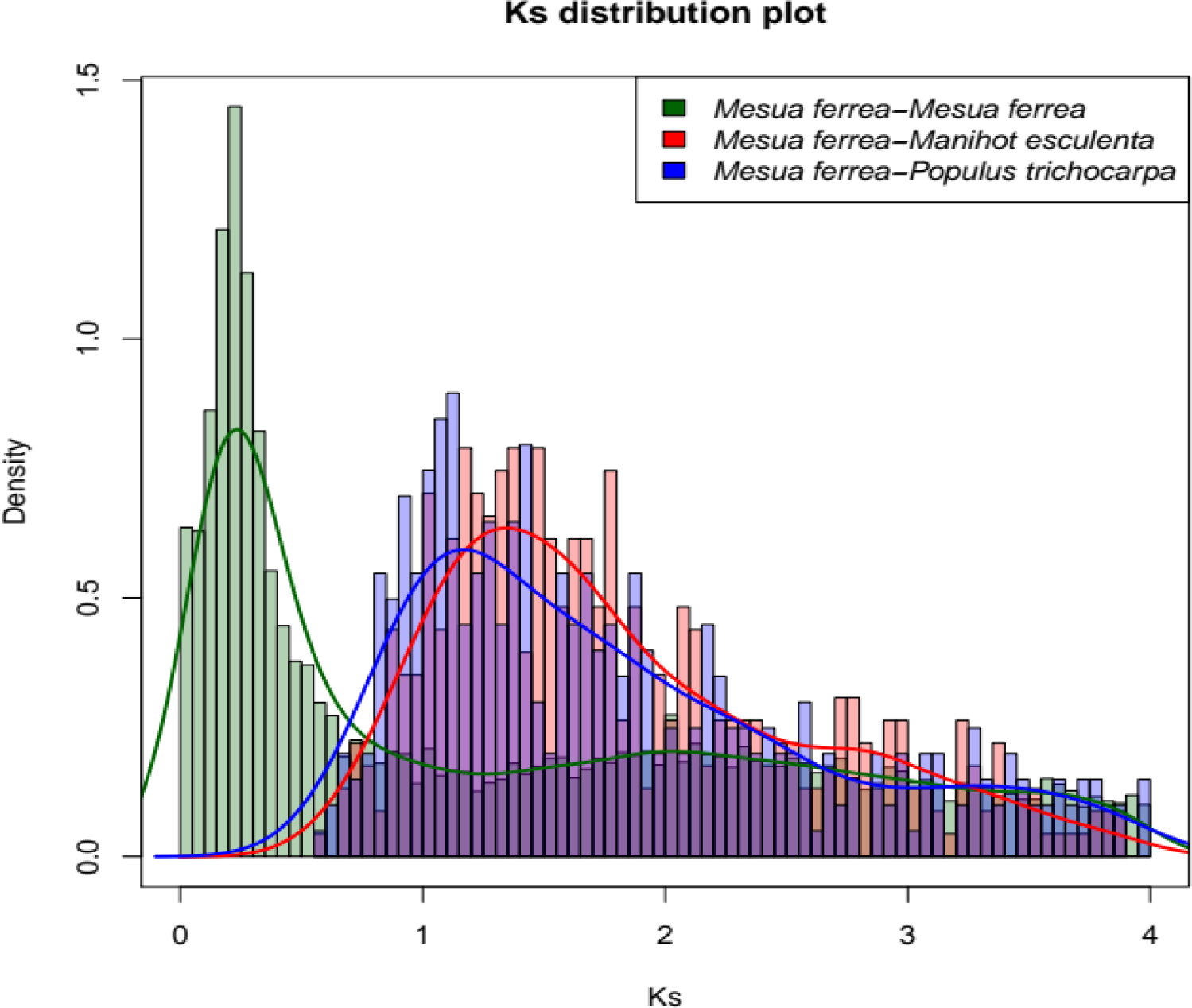
Ks-distribution plot. Distribution of synonymous substitutions (Ks) across paralogs of *Mesua ferrea* (green) and homologs of *Mesua ferrea* with *Manihot esculenta* (red) and *Populus trichocarpa* (blue). Blue and red peaks show common WGD event across Malpighiales which is around 1.1-1.2. There is a possibility of independent WGD event in clusiods or *Mesua ferrea* which shows peak at 0.2. The independent WGD event could be species-specific or clade specific but cannot be stated due to unavailability of other genomic dataset from clusioids.

**Figure S12:**
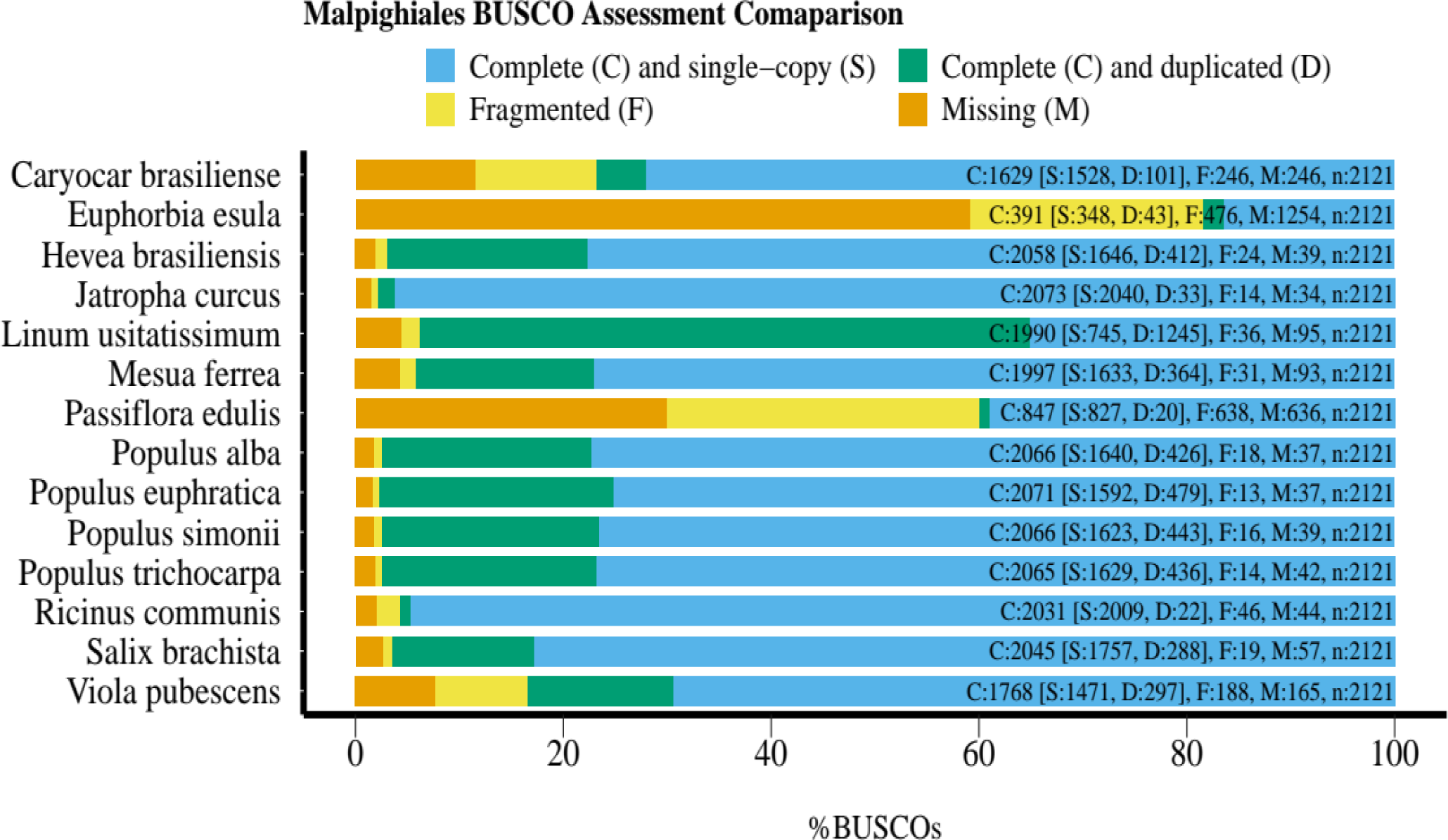
A**s**sembly **quality comparison of Malpighiales using Eudicotyledons_odb10 dataset.** Comparison of genome completeness based on BUSCO scores of Malpighiales genome assemblies. *Mesua ferrea* showed relatively complete assembly compared to other compared species.

**Figure S13a:**
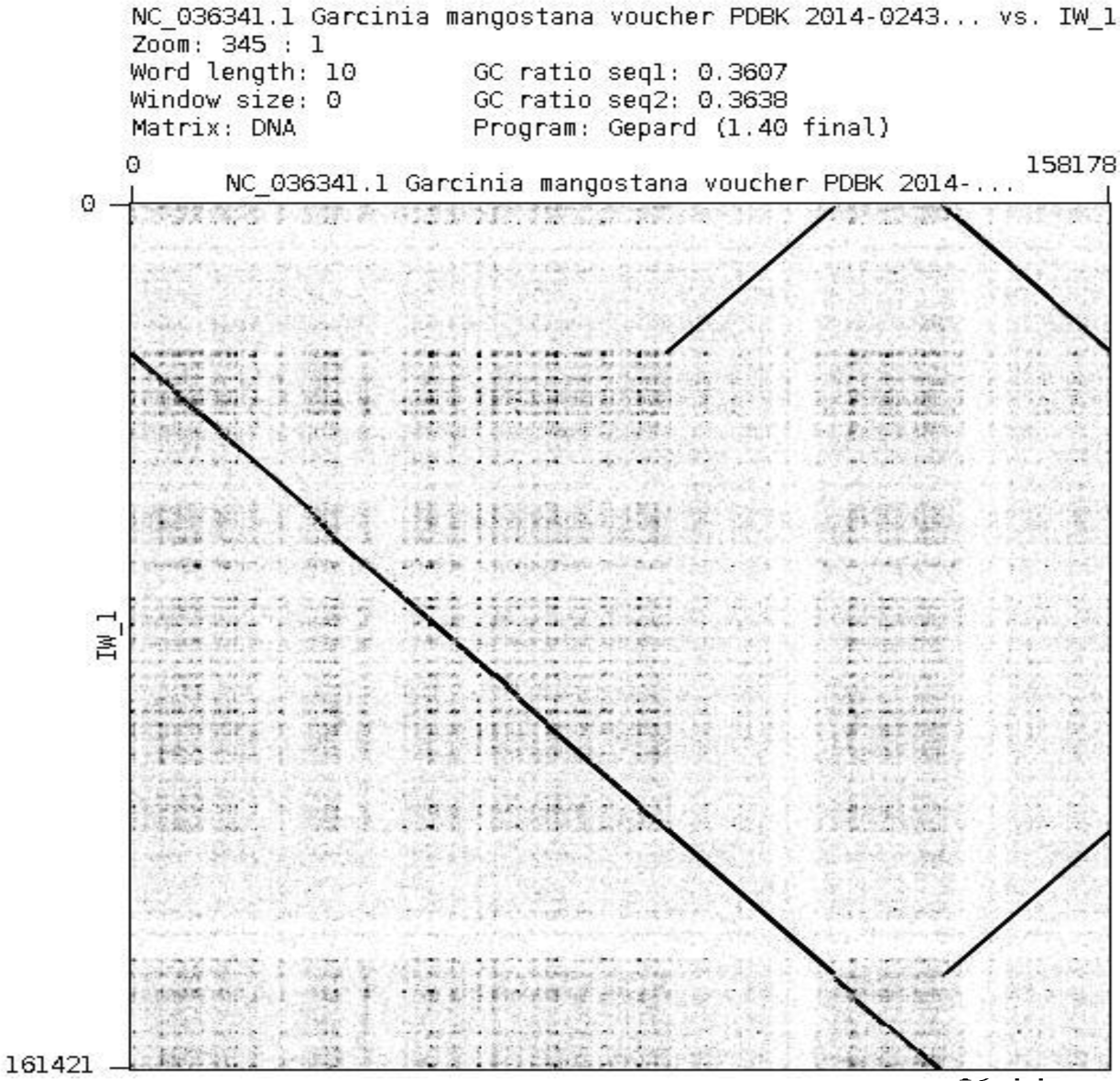
Dot-plot of *Garcinia mangostana* (top) and *Mesua ferrea* (left) plastome contig set 1.

**Figure S13b:**
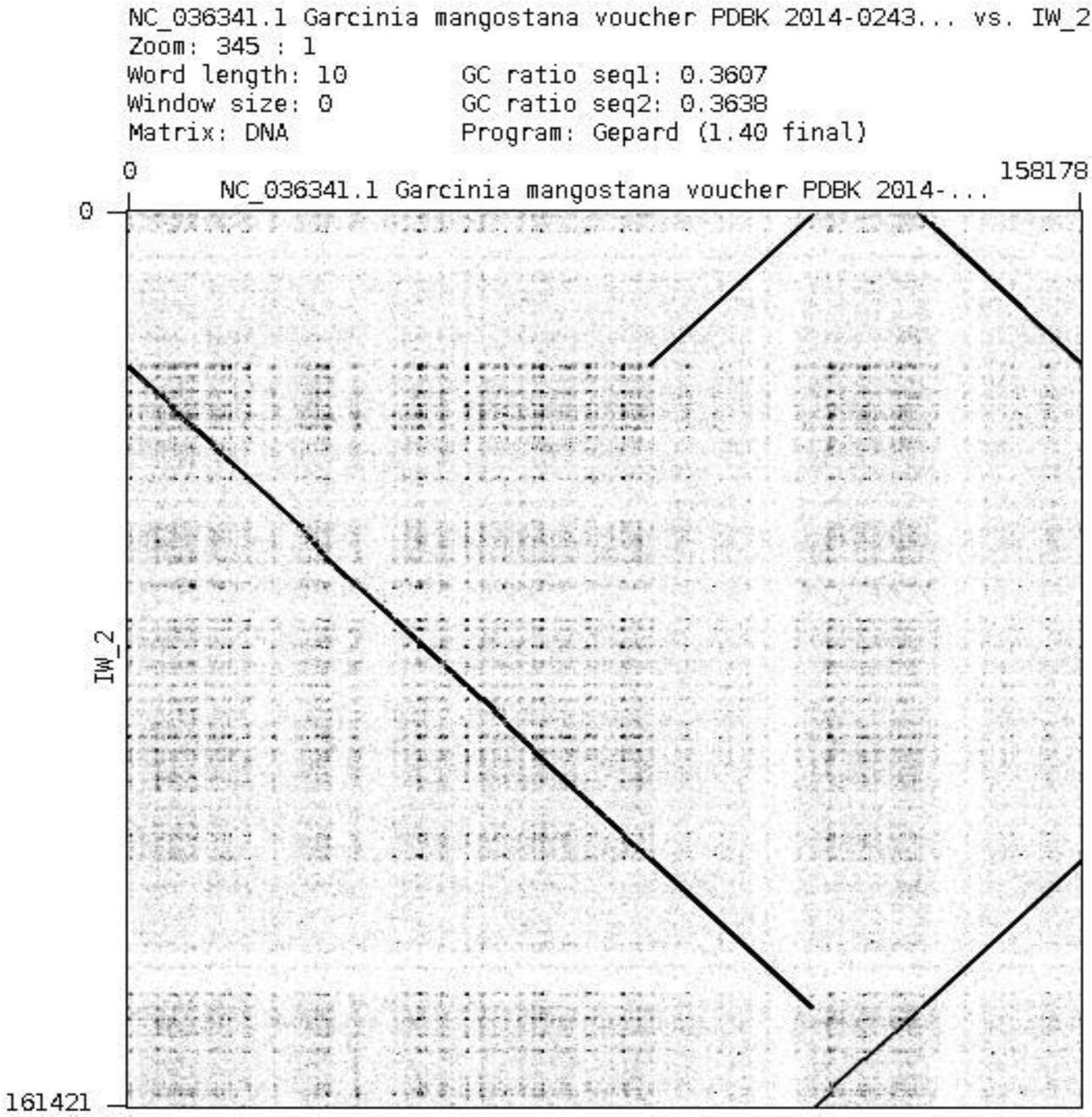
Dot-plot of *Garcinia mangostana* (top) and *Mesua ferrea* (left) plastome contig set 2.

**Figure S13c:**
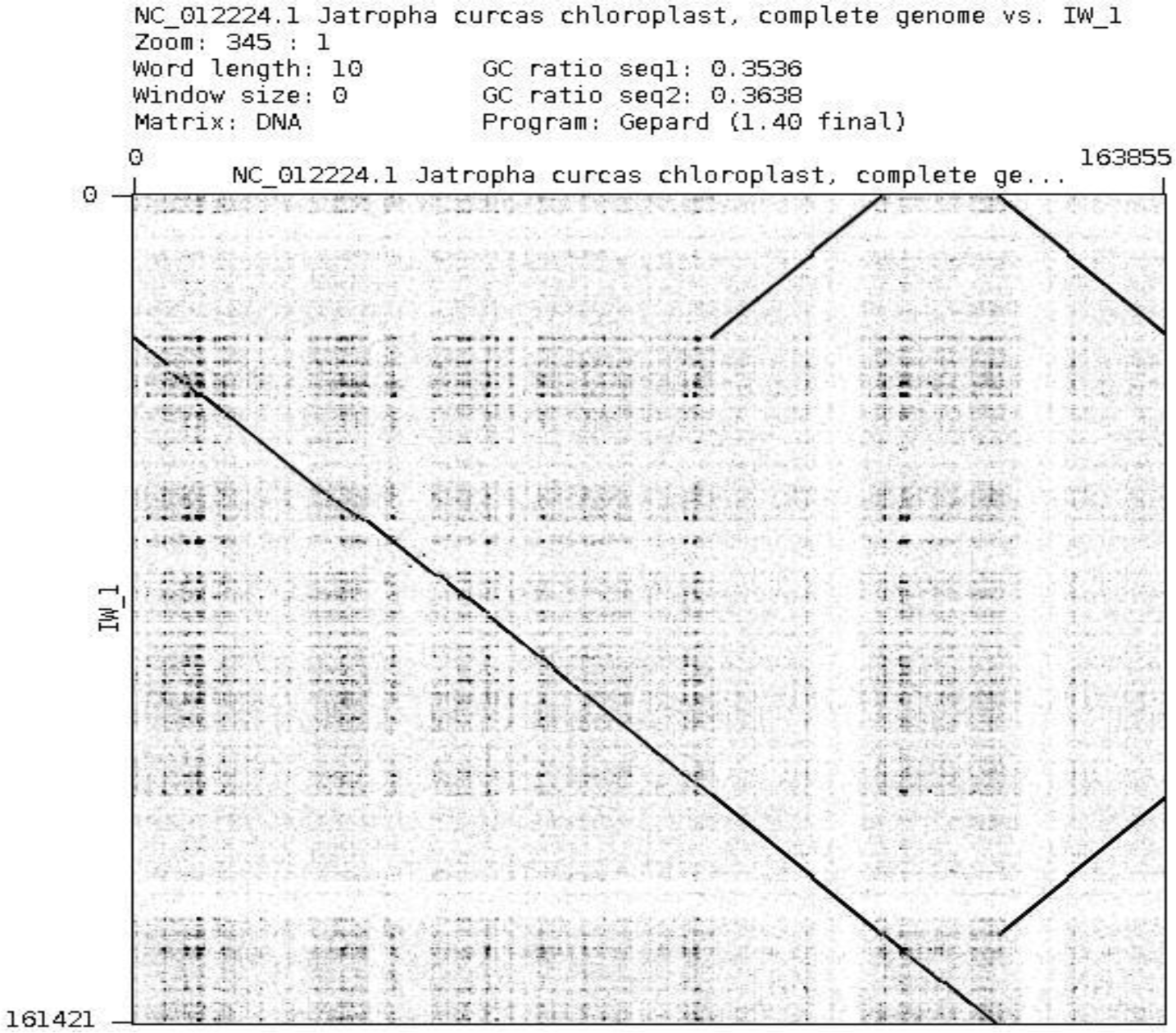
Dot-plot of *Jatropha curcus* (top) and *Mesua ferrea* (left) plastome contig set 1.

**Figure S13d:**
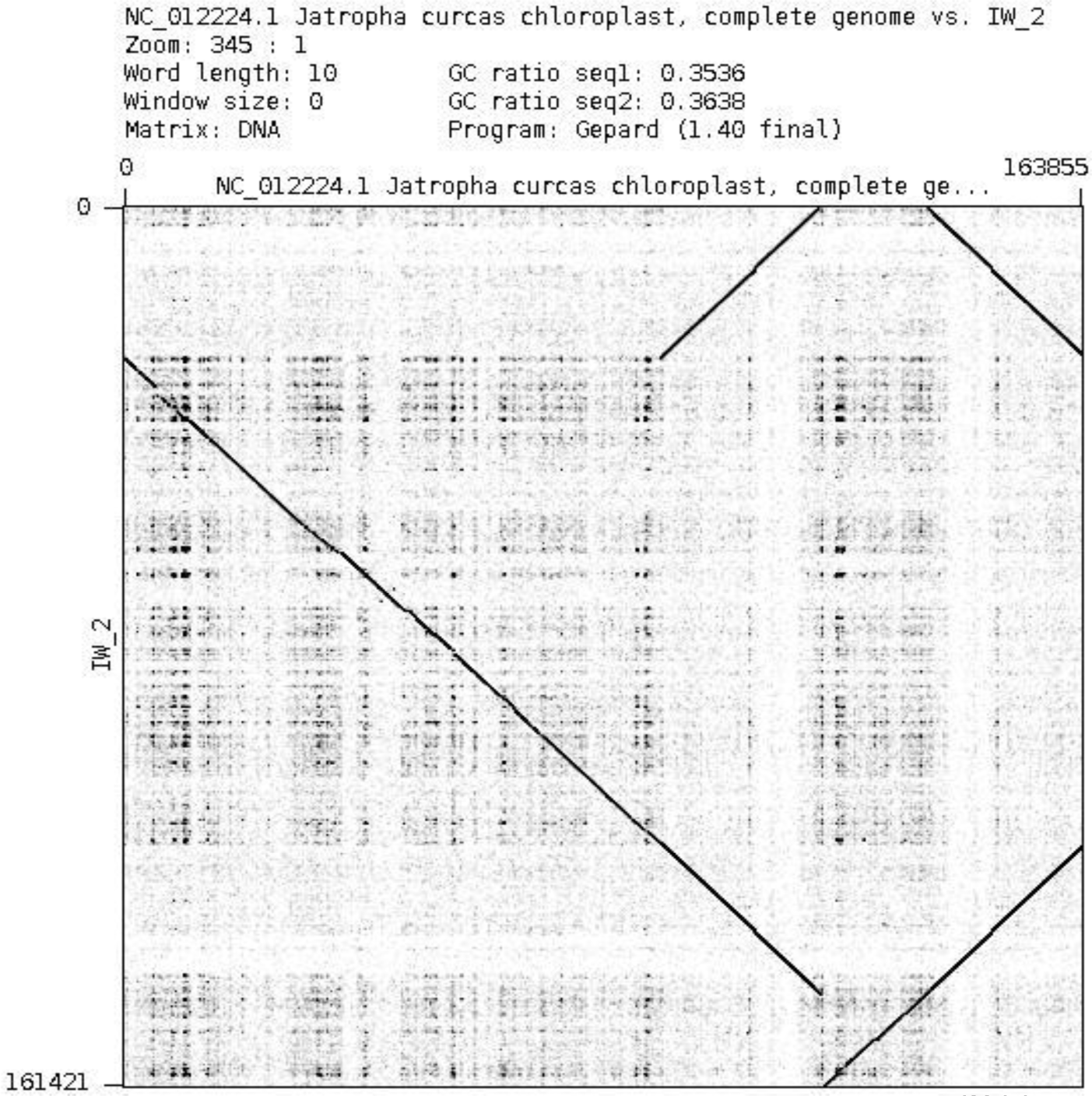
Dot-plot of *Jatropha curcus* (top) and *Mesua ferrea* (left) plastome contig set 2.

**Figure S13e:**
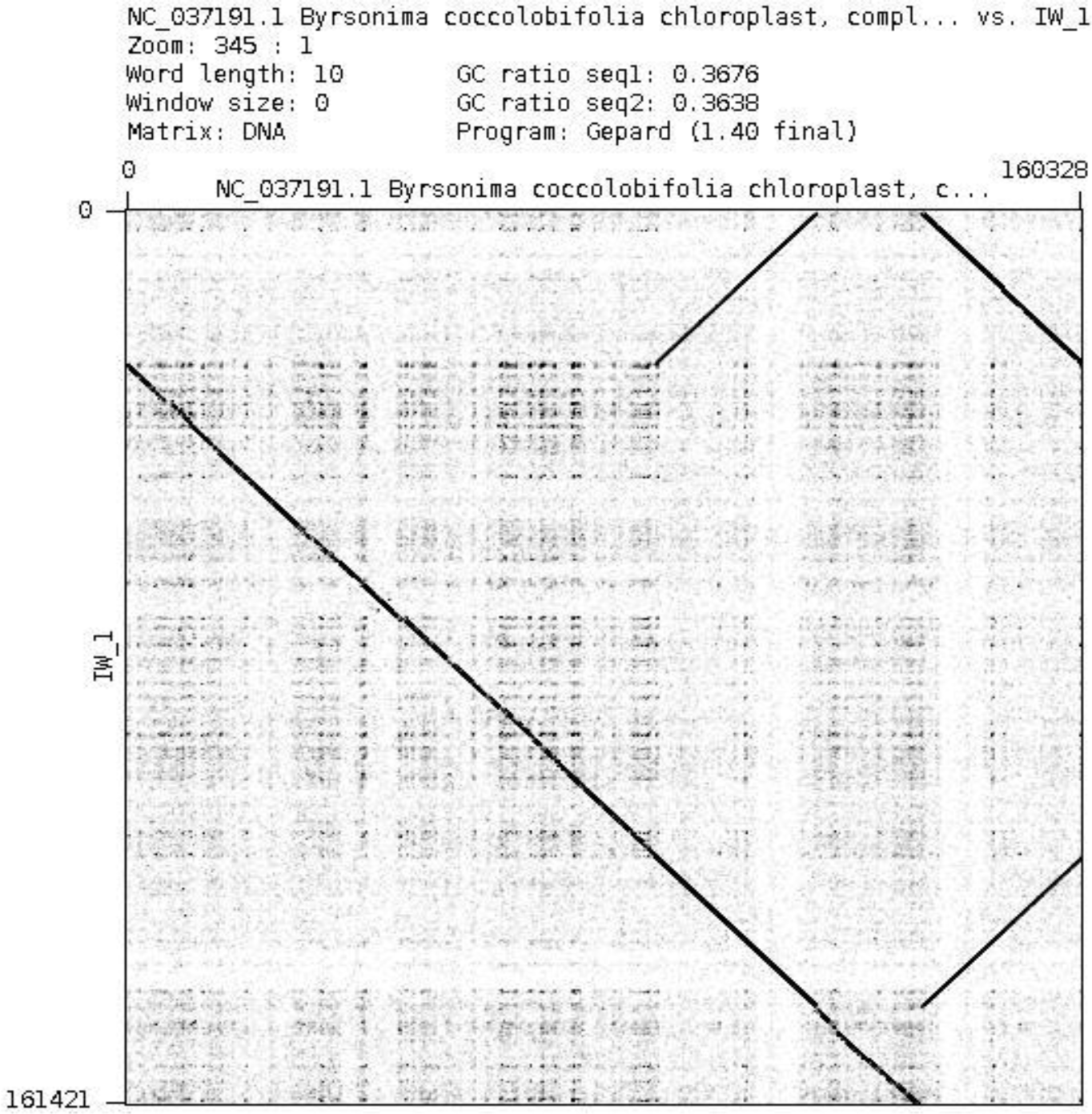
Dot-plot of *Byrsonima coccolobifolia* (top) and *Mesua ferrea* (left) plastome contig set 1.

**Figure S13f:**
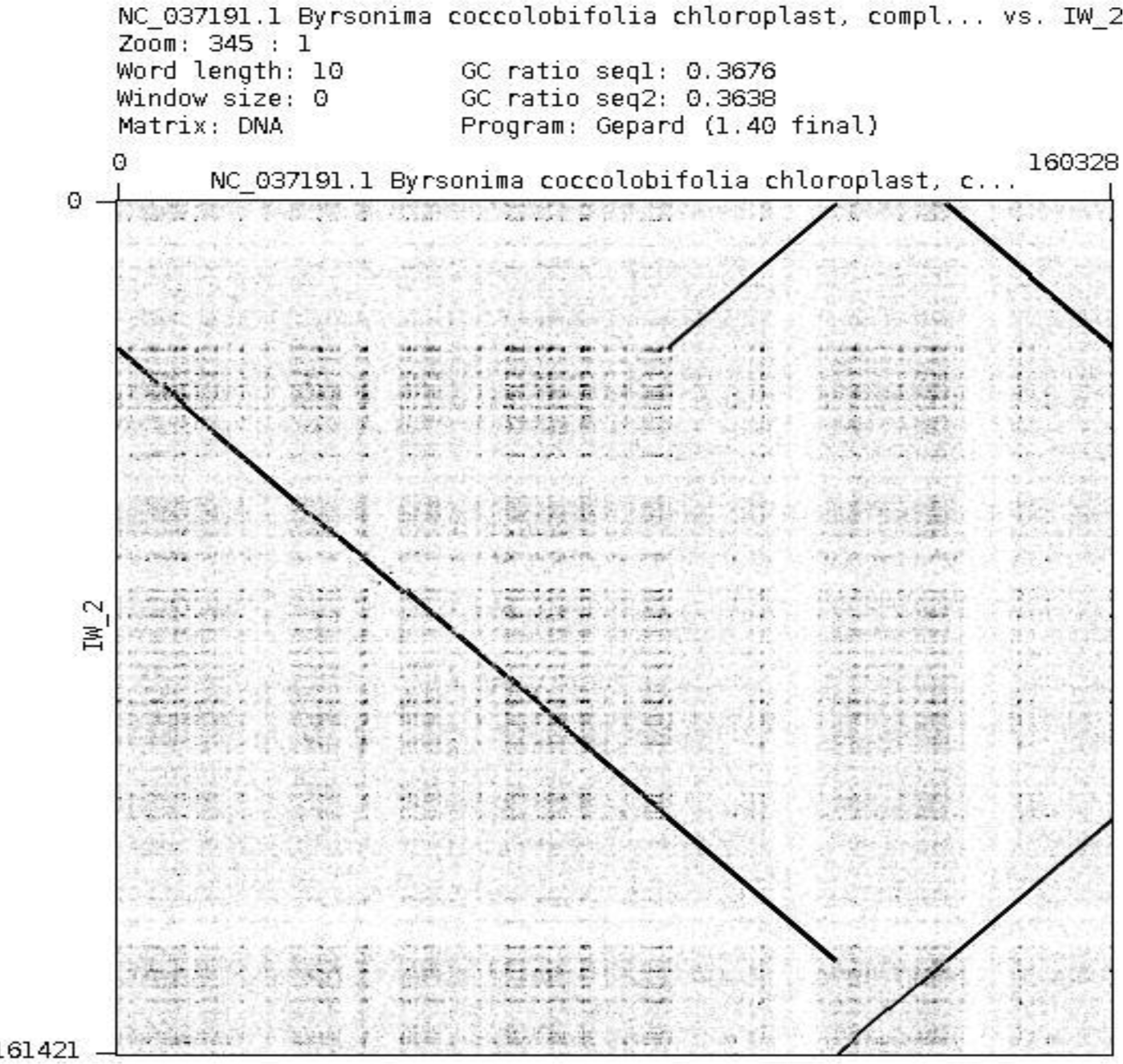
Dot-plot of *Byrsonima coccolobifolia* (top) and *Mesua ferrea* (left) plastome contig set 2.

**Figure S14:**
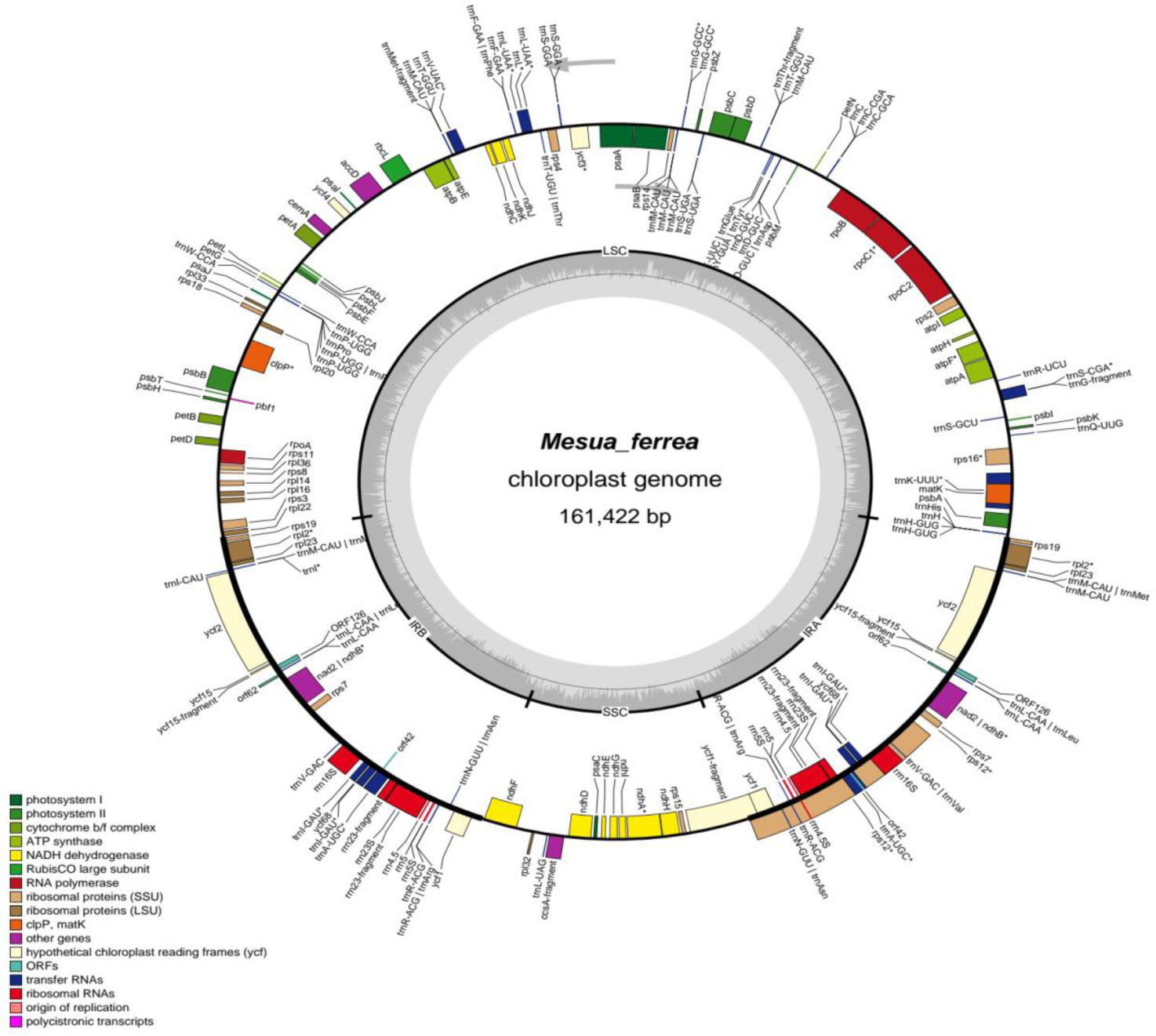
C**i**rcular **plot of *Mesua ferrea* plastome.** Assembled chloroplast genome length is 161.422 Kbp. Annotated genes are shown with colours showing their association to the pathways.

**Figure S15:**
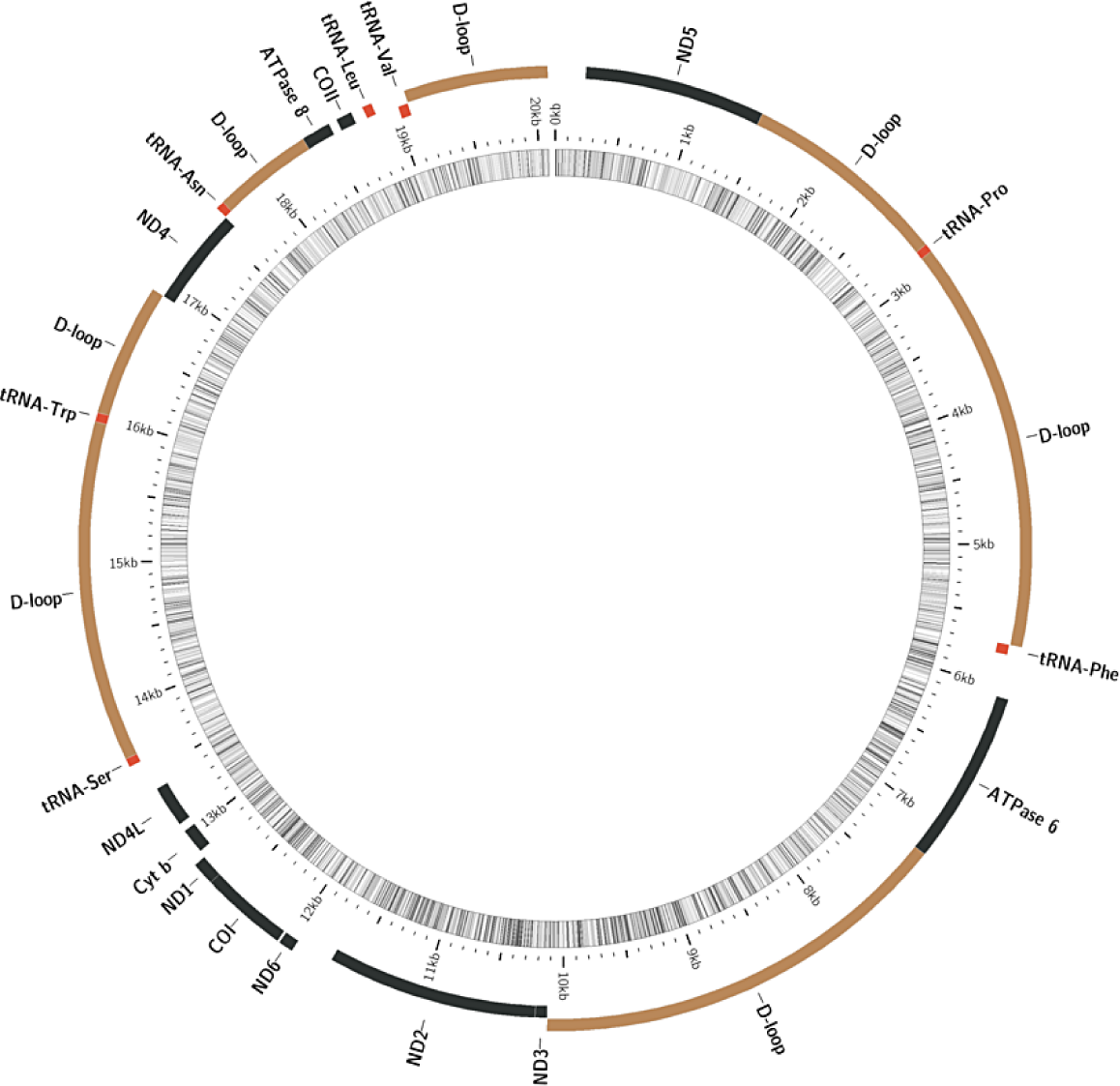
Circular plot of *Mesua ferrea* Mitochondria.

**Supplementary Table S1:**
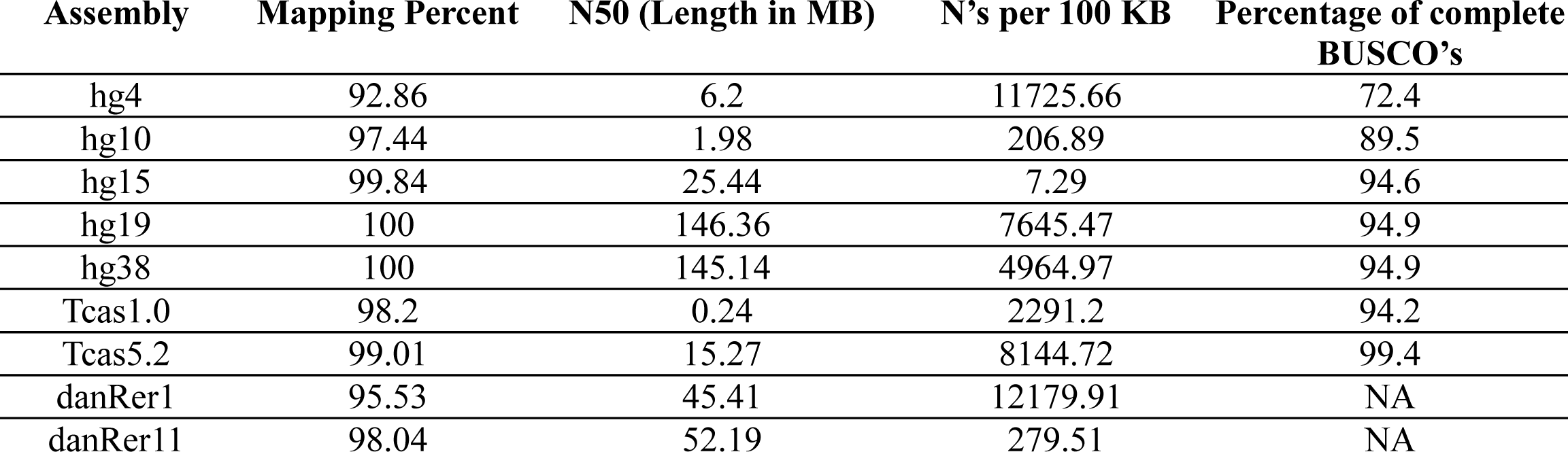
Assembly quality comparison.

**Supplementary Table S2:** SRA reads used in this study.

| Sr. No. | SRA Accession | Sample Details |
| --- | --- | --- |
| 1 | SRR9091899 | *Homo sapiens* NA12878 Illumina-Hiseq 4000 |
| 2 | SRR7121482 | *Homo sapiens* NA12878 BGISEQ-500 |
| 3 | SRR7906163 | *Populus trichocarpa* |
| 4 | SRR9007075 | *Populus euphratica* |
| 5 | SRR3045849 | *Populus nigra* |
| 6 | SRR2745904 | *Populus tremula* |
| 7 | SRR2751102 | *Populus tremuloides* |
| 8 | SRR7121482 | *Mesua ferrea* BGISEQ-500 |
| 9 | ERR2026087 | *Betula pendula* |
| 10 | SRR6058604 | *Durio zibethinus* |
| 11 | SRR10339638 | *Eucalyptus grandis* |
| 12 | SRR7072804 | *Faidherbia albida* |
| 13 | SRR5265130 | *Tectona grandis* |
| 14 | SRR3860174 | *Quercus robur* |
| 15 | SRR5678803 | *Ficus carica* |
| 16 | DRR142810 | *Citrus unshiu* |
| 17 | SRR5674478 | *Trema orientalis* |
| 18 | SRR8731963 | *Castanea mollissima* |
| 19 | ERR1346607 | *Olea europea* |
| 20 | SRR5150443 | *Santalum album* |
| 21 | SRR5019373 | *Populus pruinosa* |
| 22 | ERR3491152 | *Trochodendron aralioides* |
| 23 | SRR5992151 | *Tribolium castaneum* |
| 24 | SRR6687445 | *Danio rerio* |
| 25 | This study | *Mesua ferrea* |

**Supplementary Table S3:** Genome assembly statistics of *Mesua ferrea* genome.

| Sr. No. | Parameter | Contigs | Scaffolds |
| --- | --- | --- | --- |
| 1 | Total Assembled genome length (MB) | 609.19 | 614.35 |
| 2 | Number of contigs<sup>a</sup> | 531964 | 503130 |
| 3 | Largest contig<sup>a</sup> (KB) | 2231.61 | 4132.84 |
| 4 | N50<sup>a</sup> (KB) | 251.71 | 392.76 |
| 5 | L50<sup>a</sup> | 607 | 379 |
| 6 | L75<sup>a</sup> | 2612 | 1584 |
| 7 | GC % | 38.38 | 38.38 |

<sup>a</sup>Statistics are based on contigs > 100 bp.

**Supplementary Table S4:**
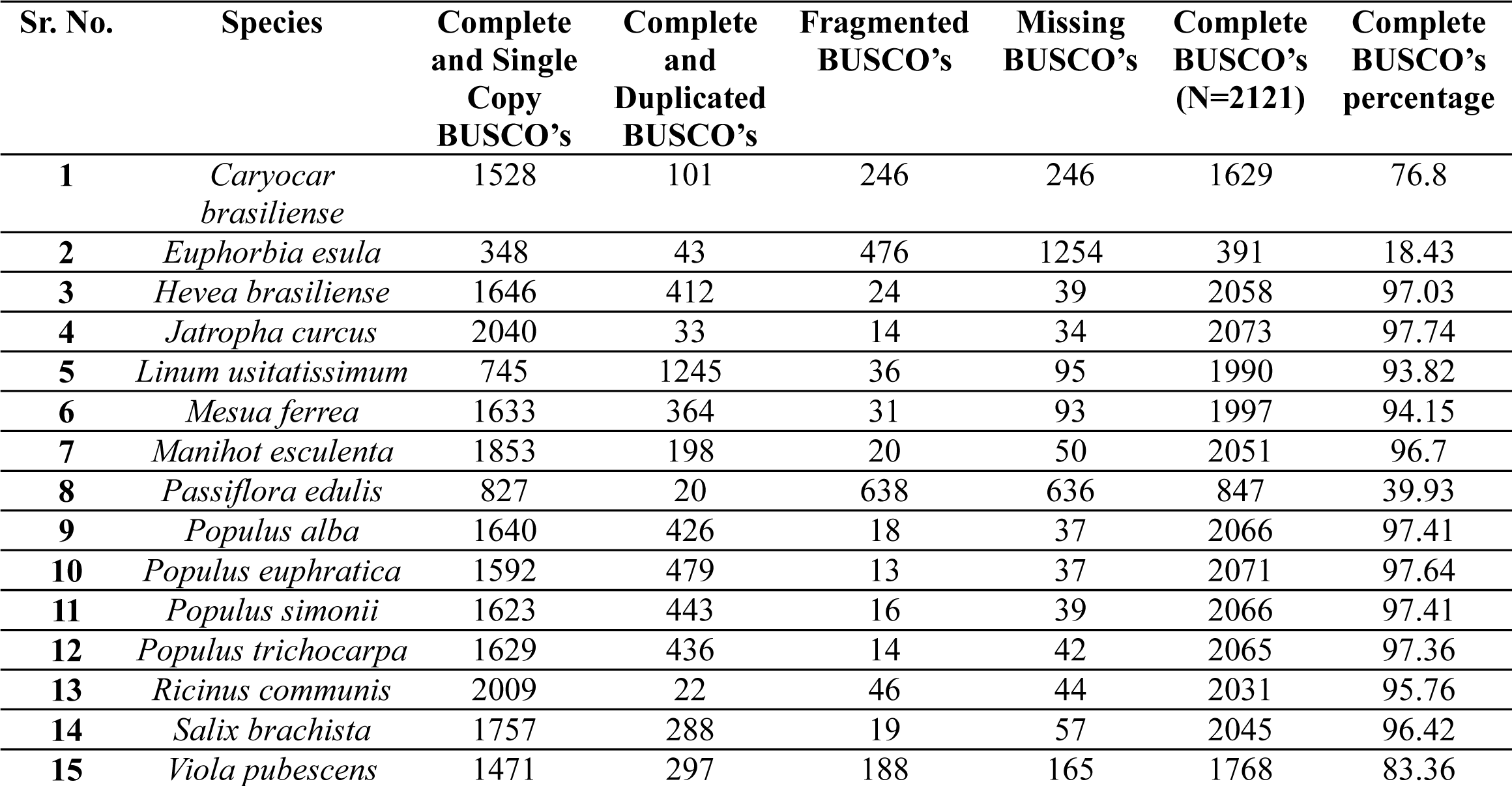
BUSCO score comparison across previously published genomes from Malpighiales using eudicotyledons_odb10 dataset.

| Sr. No. | Species | Complete and Single Copy BUSCO's | Complete and Duplicated BUSCO's | Fragmented BUSCO's | Missing BUSCO's | Complete BUSCO's (N=2121) | Complete BUSCO's percentage |
| --- | --- | --- | --- | --- | --- | --- | --- |
| 1 | *Caryocar brasiliense* | 1528 | 101 | 246 | 246 | 1629 | 76.8 |
| 2 | *Euphorbia esula* | 348 | 43 | 476 | 1254 | 391 | 18.43 |
| 3 | *Hevea brasiliense* | 1646 | 412 | 24 | 39 | 2058 | 97.03 |
| 4 | *Jatropha curcus* | 2040 | 33 | 14 | 34 | 2073 | 97.74 |
| 5 | *Linum usitatissimum* | 745 | 1245 | 36 | 95 | 1990 | 93.82 |
| 6 | *Mesua ferrea* | 1633 | 364 | 31 | 93 | 1997 | 94.15 |
| 7 | *Manihot esculenta* | 1853 | 198 | 20 | 50 | 2051 | 96.7 |
| 8 | *Passiflora edulis* | 827 | 20 | 638 | 636 | 847 | 39.93 |
| 9 | *Populus alba* | 1640 | 426 | 18 | 37 | 2066 | 97.41 |
| 10 | *Populus euphratica* | 1592 | 479 | 13 | 37 | 2071 | 97.64 |
| 11 | *Populus simonii* | 1623 | 443 | 16 | 39 | 2066 | 97.41 |
| 12 | *Populus trichocarpa* | 1629 | 436 | 14 | 42 | 2065 | 97.36 |
| 13 | *Ricinus communis* | 2009 | 22 | 46 | 44 | 2031 | 95.76 |
| 14 | *Salix brachista* | 1757 | 288 | 19 | 57 | 2045 | 96.42 |
| 15 | *Viola pubescens* | 1471 | 297 | 188 | 165 | 1768 | 83.36 |

**Supplementary Table S5:** BUSCO score comparison across previously published genomes from Malpighiales using embryophyta_odb10 dataset.

| Sr. No. | Species | Complete and Single Copy BUSCO's | Complete and Duplicated BUSCO's | Fragmented BUSCO's | Missing BUSCO's | Complete BUSCO's (N=1375) | Complete BUSCO's percentage |
| --- | --- | --- | --- | --- | --- | --- | --- |
| 1 | *Caryocar brasiliense* | 1017 | 43 | 183 | 132 | 1060 | 77.1 |
| 2 | *Euphorbia esula* | 265 | 42 | 424 | 644 | 307 | 22.4 |
| 3 | *Hevea brasiliense* | 1088 | 243 | 16 | 28 | 1331 | 96.8 |
| 4 | *Jatropha curcas* | 1331 | 18 | 4 | 22 | 1349 | 98.1 |
| 5 | *Linum usitatissimum* | 467 | 859 | 10 | 39 | 1326 | 96.5 |
| 6 | *Mesua ferrea* | 1093 | 199 | 18 | 65 | 1292 | 94 |
| 7 | *Manihot esculenta* | 1248 | 80 | 15 | 32 | 1328 | 96.6 |
| 8 | *Passiflora edulis* | 598 | 9 | 460 | 308 | 607 | 44.2 |
| 9 | *Populus alba* | 1074 | 270 | 10 | 21 | 1344 | 97.7 |
| 10 | *Populus euphratica* | 1063 | 284 | 6 | 22 | 1347 | 98 |
| 11 | *Populus simonii* | 1076 | 270 | 8 | 21 | 1346 | 97.9 |
| 12 | *Populus trichocarpa* | 1084 | 258 | 9 | 24 | 1342 | 97.6 |
| 13 | *Ricinus communis* | 1317 | 7 | 19 | 32 | 1375 | 96.3 |
| 14 | *Salix brachista* | 1200 | 146 | 5 | 24 | 1346 | 97.9 |
| 15 | *Viola pubescens* | 973 | 186 | 124 | 92 | 1159 | 84.3 |

**Supplementary Table S6:** LTR-retriever LAI scores for Malpighiales genome assemblies.

| Sr. No. | Species | Raw LAI | LAI |
| --- | --- | --- | --- |
| 1 | *Hevea brasiliensis* | 1.75 | 0.75 |
| 2 | *Jatropha curcas* | 2.37 | 1.22 |
| 3 | *Linum usitatissimum* | 5.55 | 8.76 |
| 4 | *Manihot esculenta* | 3.79 | 3.5 |
| 5 | *Mesua ferrea* | 2.1 | 7.73 |
| 6 | *Populus alba* | 11.32 | 11.74 |
| 7 | *Populus euphratica* | 1.25 | 3 |
| 8 | *Populus simonii* | 8.96 | 12.15 |
| 9 | *Populus trichocarpa* | 6.54 | 11.57 |
| 10 | *Rhizophora apiculata* | 7.48 | 13.03 |
| 11 | *Ricinus communis* | 4.37 | 5.43 |
| 12 | *Salix brachista* | 17.54 | 17.1 |

**Supplementary Table S7:** Annotation Statistics for iterative MAKER-P annotation of *Mesua ferrea* genome assembly.

| Parameter | Round 1 | Round 2 | Round 3 | Round 4 | Round 5 |
| --- | --- | --- | --- | --- | --- |
| No. of protein coding genes | 35557 | 38971 | 38965 | 39011 | 46540 |
| Gene density per KB | 0.07 | 0.06 | 0.08 | 0.06 | 0.08 |
| Average gene length (bp) | 2631.77 | 2722.22 | 2768.24 | 2761.98 | 2477.54 |
| Average exons per mRNA | 4.73 | 4.96 | 5.1 | 5.09 | 4.55 |
| Average exon length (bp) | 206.42 | 217.37 | 222.09 | 222.14 | 222.24 |
| Average intron length (bp) | 401.68 | 378.83 | 359.1 | 360.17 | 365.12 |
| Cumulative fraction of genes with AED < 0.5 | 0.99 | 0.92 | 0.92 | 0.92 | 0.93 |
| Percentage of complete BUSCO's | 89.4 | 93.3 | 92.3 | 92.1 | 94.7 |

**Supplementary Table S8:** Genome assemblies used in this study.

| Sr. No. | Species | Assembly | Database |
| --- | --- | --- | --- |
| 1 | *Homo sapiens* | GRCh38.p13 | ENSEMBL |
| 2 | *Homo sapiens* | hg4 | UCSC Genome Browser |
| 3 | *Homo sapiens* | hg10 | UCSC Genome Browser |
| 4 | *Homo sapiens* | hg15 | UCSC Genome Browser |
| 5 | *Homo sapiens* | hg19 | UCSC Genome Browser |
| 6 | *Caryocar brasiliense* | Cbr_v1.0 | NCBI |
| 7 | *Euphorbia esula* | ASM291907v1 | NCBI |
| 8 | *Hevea brasiliense* | ASM165405v1 | NCBI |
| 9 | *Jatropha curcus* | JatCur_1.0 | NCBI |
| 10 | *Linum usitatissimum* | ASM22429v2 | NCBI |
| 11 | *Mesua ferrea* | This Study |  |
| 12 | *Manihot esculenta* | Manihot esculenta v6 | NCBI |
| 13 | *Passiflora edulis* | ASM215610v1 | NCBI |
| 14 | *Populus alba* | ASM523922v1 | NCBI |
| 15 | *Populus euphratica* | PopEup_1.0 | NCBI |
| 16 | *Populus simonii* | Populus simonii_2.0 | NCBI |
| 17 | *Populus trichocarpa* | Pop_tri_v3 | NCBI |
| 18 | *Ricinus communis* | JCVI_RCG_1.1 | NCBI |
| 19 | *Salix brachista* | ASM907833v1 | NCBI |
| 20 | *Viola pubescens* | GCA_002752925.1 | NCBI |
| 21 | *Betula pendula* | Bpev01 | NCBI |
| 22 | *Durio zibethinus* | Duzib1.0 | NCBI |
| 23 | *Eucalyptus grandis* | Egrandis1_0 | NCBI |
| 24 | *Faidherbia albida* | http://gigadb.org/dataset/101054 | GIGADB |
| 25 | *Tectona grandis* | https://biit.cs.ut.ee/supplementary/WGSteak/ |  |
| 26 | *Quercus robur* | Q_robur_v1 | NCBI |
| 27 | *Ficus carica* | UNIPi_FiCari_1.0 | NCBI |
| 28 | *Citrus unshiu* | http://www.citrusgenome.jp/ | DDBJ |
| 29 | *Trema orientalis* | TorRG33x02_asm01 | NCBI |
| 30 | *Castanea mollissima* | ASM76360v1 | NCBI |
| 31 | *Olea europea* | O_europea_v1 | NCBI |
| 32 | *Santalum album* | ASM291163v1 | NCBI |
| 33 | *Populus pruinosa* | http://gigadb.org/dataset/100319 | GIGADB |
| 34 | *Trochodendron aralioides* | http://gigadb.org/dataset/100657 | GIGADB |
| 35 | *Tribolium castaneum* | ftp://ftp.hgsc.bcm.edu/Tcastaneum/Tcas1.0/ | HGEC, BGM |
| 36 | *Tribolium castaneum* | Tcas5.2 | NCBI |
| 37 | *Danio rerio* | danRer1 | UCSC Genome Browser |
| 38 | *Danio rerio* | danRer11 | UCSC Genome Browser |

**Supplementary Table S9:** Details of the mutation rate and generation time used. The Median estimate of mutation rate i.e. 2.5e-09 per site per year was used for all and converted to per generation mutation rates according to their generation times.

| Sr. No. | Species | Generation time used | Mutation rate per generation |
| --- | --- | --- | --- |
| 1 | *Betula pendula* | 20 | 5e-08 |
| 2 | *Durio zibethinus* | 7 | 1.75e-08 |
| 3 | *Eucalyptus grandis* | 7 | 1.75e-08 |
| 4 | *Faidherbia albida* | 7 | 1.75e-08 |
| 5 | *Tectona grandis* | 7 | 1.75e-08 |
| 6 | *Quercus robur* | 15 | 3.75e-08 |
| 7 | *Ficus carica* | 7 | 1.75e-08 |
| 8 | *Citrus unshiu* | 7 | 1.75e-08 |
| 9 | *Trema orientalis* | 7 | 1.75e-08 |
| 10 | *Castanea mollissima* | 7 | 1.75e-08 |
| 11 | *Olea europea* | 7 | 1.75e-08 |
| 12 | *Santalum album* | 7 | 1.75e-08 |
| 13 | *Populus pruinosa* | 15 | 3.75e-08 |
| 14 | *Trochodendron araloides* | 15 | 3.75e-08 |
| 15 | *Mesua ferrea* | 15 | 3.75e-08 |
| 16 | *Tribolium castaneum* | 0.3 (12 weeks) | 2.7e-09 |
| 17 | *Danio rerio* | 1 | 1e-09 |
| 18 | *Homo sapiens* | 25 | 2.5e-08 |

